# Interneuron subtypes enable independent modulation of excitatory and inhibitory firing rates after sensory deprivation

**DOI:** 10.1101/2021.05.25.445562

**Authors:** Leonidas M. A. Richter, Julijana Gjorgjieva

## Abstract

Diverse interneuron subtypes determine how cortical circuits process sensory information depending on their connectivity. Sensory deprivation experiments are ideally suited to unravel the plasticity mechanisms which shape circuit connectivity, but have yet to consider the role of different inhibitory subtypes. We investigate how synaptic changes due to monocular deprivation affect the firing rate dynamics in a microcircuit network model of the visual cortex. We demonstrate that, in highly recurrent networks, deprivation-induced plasticity generates fundamentally different activity changes depending on interneuron composition. Considering parvalbumin-positive (*PV*^+^) and somatostatin-positive (*SST*^+^) interneuron subtypes can capture the experimentally observed independent modulation of excitatory and inhibitory activity during sensory deprivation when *SST*^+^ feedback is sufficiently strong. Our model also applies to whisker deprivation in the somatosensory cortex revealing that these mechanisms are general across sensory cortices. Therefore, we provide a mechanistic explanation for the differential role of interneuron subtypes in regulating cortical dynamics during deprivation-induced plasticity.

## Introduction

Cortical interneurons are a diverse set of cell types that serve multiple cell-type-specific functions in neural computations (Tremblay et al., 2016; Wood et al., 2017). The connectivity among excitatory pyramidal neurons and different subtypes of interneurons in feedforward and recurrent neural circuits plays a key role in establishing these functions. In mature cortical circuits, these interneuron subtypes are involved in disinhibition during locomotion and learning (Abs et al., 2018; Adler et al., 2019; Fu et al., 2014; Letzkus et al., 2011; Niell and Stryker, 2010), response reversal during top-down modulation (Dipoppa et al., 2018; Fu et al., 2014; Garcia del Molino et al., 2017; Keller et al., 2020; Pakan et al., 2016), and shown to affect surround suppression (Adesnik et al., 2012; Ayaz et al., 2013; Litwin-Kumar et al., 2016; Ozeki et al., 2009) and excitatory tuning (Atallah et al., 2012; Lee et al., 2012; Wilson et al., 2018). Inhibitory synapses are plastic (D’amour and Froemke, 2015; Field et al., 2020); however, we still do not understand how the plasticity of connections among the different interneuron subtypes and excitatory neurons shapes circuit dynamics and computations.

Synaptic plasticity in cortical circuits is particularly prominent in development and young adulthood during so-called *critical periods* (Hensch, 2005). Experiments have used the sensitivity of cortical circuits to perturbations during the critical period to unravel the plasticity mechanisms which shape circuit connectivity and resulting activity (Espinosa and Stryker, 2012; Gainey and Feldman, 2017; Turrigiano, 2011; Wiesel, 1982). An important paradigm to investigate these mechanisms is monocular deprivation (MD), the sustained closure of one eye (Kuhlman et al., 2013; Maffei et al., 2006, 2004; Miska et al., 2018; Wang et al., 2013; Wiesel and Hubel, 1963). This sustained closure causes a biphasic response in the monocular region of primary visual cortex (V1m), driven exclusively by the contralateral eye, that first reduces and then restores excitability (Hengen et al., 2013, 2016; Kaneko et al., 2008). The plasticity of inhibitory synapses has been shown to regulate these processes (Kannan et al., 2016; Maffei et al., 2010, 2006; Miska et al., 2018; Nahmani and Turrigiano, 2014). However, it primarily pertains to fast-spiking interneurons, which most likely correspond to parvalbumin-positive (*PV*^+^) interneurons, the most abundant and best-studied subtype of interneurons in the cortex (Hu et al., 2014; Tremblay et al., 2016).

Experimental and modeling work has found that the plasticity of recurrent connectivity, and especially the potentiation of intracortical inhibition, dominates over the depression of feedforward connectivity to explain the initial decrease of excitatory and inhibitory activity after MD (Miska et al., 2018). In turn, homeostatic mechanisms involving both excitatory and inhibitory circuit components recover different aspects of network dynamics after deprivation (Ma et al., 2019; Wu et al., 2020). However, recent experiments show that the network behavior might be more complex. Namely, while inhibitory neurons decrease their firing rates one day after MD induction, excitatory neurons are delayed by an additional day. What mechanism lies behind this independent modulation of excitatory and inhibitory firing rates remains unclear.

We used a recurrent network with balanced excitation and inhibition to study this process in a microcircuit model of the visual cortex. Theoretical work has shown that the dynamics and computational properties of these networks are highly dependent on the network’s operating regime, which is determined by the strength of recurrent coupling (Brunel, 2000; Hennequin et al., 2018; Murphy and Miller, 2009; Renart et al., 2010; van Vreeswijk and Sompolinsky, 1996). Although more relevant experimentally, strong excitatory recurrent coupling needs to be stabilized by sufficiently strong inhibition giving rise to *inhibition stabilized networks* (ISNs) (Ozeki et al., 2009; Tsodyks et al., 1997). A strong signature of inhibition stabilization is the *paradoxical effect,* which refers to the decrease of inhibitory firing rate in response to direct excitatory drive to inhibitory interneurons (Tsodyks et al., 1997). Recent experiments have confirmed the presence of the paradoxical effect in cortical circuits, suggesting they operate in the ISN regime (Mahrach et al., 2020; Sanzeni et al., 2020). This raises the important question of whether ISNs can explain the independent modulation of excitatory and inhibitory firing rates after brief MD.

We found that, due to the paradoxical effect, ISNs cannot capture the independent modulation of excitatory and inhibitory firing rates after brief MD. Inspired by theoretical work that generalizes the paradoxical effect to networks with multiple inhibitory interneuron subtypes (Garcia del Molino et al., 2017; Litwin-Kumar et al., 2016; Mahrach et al., 2020), we also modeled somatostatin-positive *(SST*^+^) interneurons, the second most abundant subtype of interneurons in the cortex (Tremblay et al., 2016). Our results demonstrate that the addition of *SST*^+^-interneurons inverts the firing rate response of *PV*^+^-interneurons relative to excitatory neurons in response to MD-induced plasticity by reversing the paradoxical effect. A key parameter for this effect is the strength of the feedback from *SST*^+^-interneurons to *PV*^+^-interneurons and excitatory neurons. Hence, our finding explains the independent modulation of excitatory and inhibitory firing rates, consistent with their sequential suppression during early MD with inhibitory preceding excitatory firing rates (Hengen et al., 2013, 2016). We also found similar activity changes following whisker deprivation in the somatosensory cortex, which affects interneuron intrinsic excitability rather than synaptic strength onto interneurons, suggesting that similar principles might be at work in different sensory cortices. Therefore, our work provides a mechanistic explanation for the experimentally observed temporal modulation of excitatory and inhibitory activity after sensory deprivation-induced plasticity. It also establishes a more general framework to study how the interaction of three factors – cortical operating regime, interneuron diversity, and deprivation-induced plasticity – shapes circuit dynamics and computations.

## Results

### Changes in network firing rates from MD-induced plasticity

To investigate excitatory and inhibitory activity changes in response to synaptic plasticity after brief monocular deprivation (MD), we built a network model of the primary visual cortex consisting of excitatory and inhibitory spiking neurons (Fig. 1A and Methods). Following most previous theoretical work, we initially only modeled a single class of interneurons, which we equated with parvalbumin-positive *(PV*^+^), the largest class of interneurons in the mammalian neocortex (Hu et al., 2014; Pfeffer et al., 2013; Tremblay et al., 2016). The networks have strong recurrent coupling, denoted by the overall coupling scale *J*. The coupling scale describes the operating state of the network (Lien and Scanziani, 2013; Ozeki et al., 2009; Sanzeni et al., 2020) and determines the networks’ dynamical and computational properties (Brunel, 2000; Hennequin et al., 2018; Murphy and Miller, 2009; Renart et al., 2010; van Vreeswijk and Sompolinsky, 1996). This strong recurrent coupling, however, needs to be stabilized by sufficiently strong inhibition denoted by the scaling parameter *g_rc_* (Fig. 1A) (Ozeki et al., 2009; Tsodyks et al., 1997). Such an inhibition-dominated network produces sparse, asynchronous, and irregular spiking activity (Brunel, 2000; van Vreeswijk and Sompolinsky, 1996; Wolf et al., 2014).

**Figure 1:**
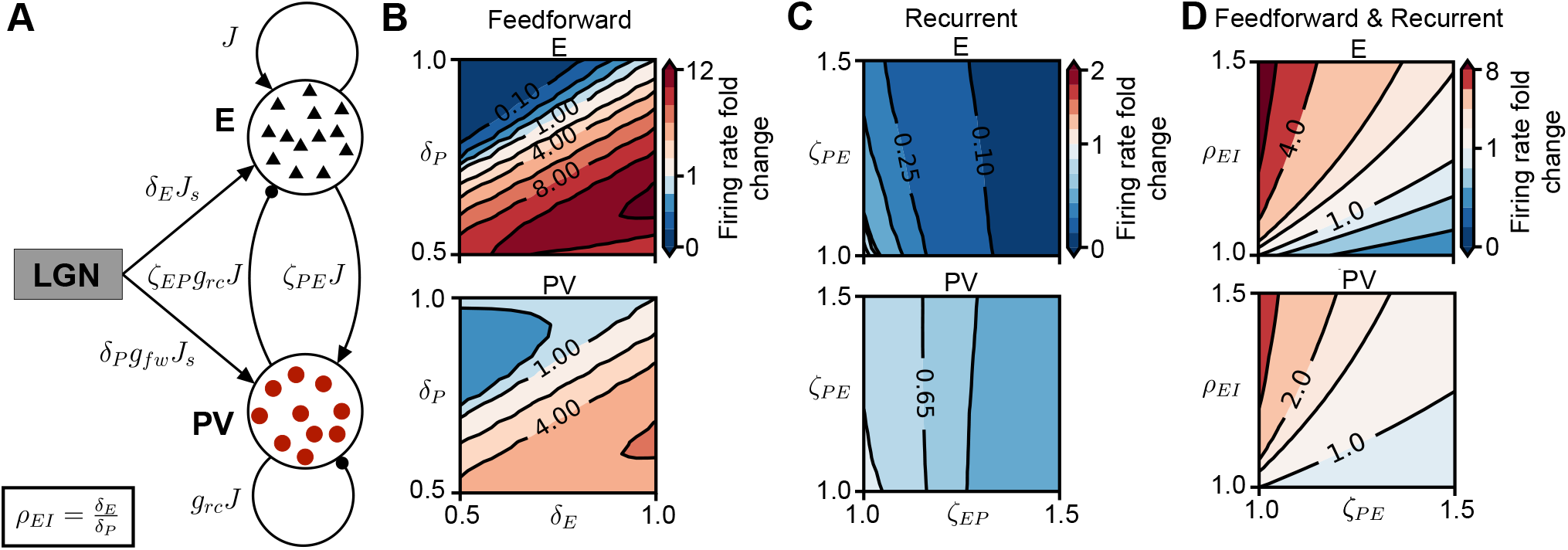
Spiking network response to synaptic changes induced by brief MD. **(A)** Network schematic showing synaptic connections among neurons with *J* denoting the overall coupling scale, *g_rc_* the dominance of recurrent inhibition and *g_fw_* the dominance of feedforward inhibition. The parameters to model MD-induced synaptic plasticity are: depression of feedforward drive to excitatory neurons *(δ_E_* < 1) and to *PV*^+^-interneurons (*δ_P_* < 1), and potentiation of recurrent excitation to *PV*^+^-interneurons (*ζ_PE_* > 1) and of recurrent inhibition from *PV*^+^ to excitatory neurons (*ζ_EP_* > 1). Values of all parameters are provided in Table 1. **(B)** Network firing rate in the (*δ_E_, δ_P_*) plane as fold-change of baseline firing rate (top right corner where *δ_E_ = δ_P_* = 1) for excitatory neurons (top) and *PV*^+^-interneurons (bottom). **(C)** Network firing rate in the *(ζ_EP_*, *ζ_PE_*) plane as fold-change of baseline firing rate (bottom left corner where *ζ_EP_ = ζ_PE_* = 1). **(D)** Combined feedforward (through the E/I ratio of feedforward synaptic changes, *ρ_EI_ = δ_E_/δ_P_*) and recurrent plasticity (through the potentiation of recurrent excitation to *PV*^+^-interneurons, *ζ_PE_*). Network firing rate in the *(ζ_PE_*, *ρ_EI_*) plane as fold-change of baseline firing rate (bottom left corner where *ζ_PE_* = *ρ_EI_* = 1).

We first modeled synaptic changes measured experimentally in the cortical circuits during brief (1-2 days of) MD, both along the feedforward pathway from the LGN to the cortex, and in the recurrent cortical circuit. In particular, feedforward excitatory synaptic inputs from the LGN to both excitatory and inhibitory neurons in the cortex depress during MD (Lambo and Turrigiano, 2013; Smith et al., 2009), while recurrent intracortical inhibition potentiates (Kannan et al., 2016; Maffei et al., 2010, 2006; Miska et al., 2018; Nahmani and Turrigiano, 2014) (but see Khibnik et al. (2010)). To investigate the relative contribution of these two factors – decrease in feedforward excitation vs. increase in recurrent inhibition – we introduced four model parameters to represent the synaptic changes (Maffei et al., 2006; Miska et al., 2018) (Fig. 1A, Table 1): (1) depression of feedforward synaptic strengths to excitatory neurons (*δ_E_* < 1), (2) depression of feedforward synaptic strengths to *PV*^+^-interneurons (*δ_P_* < 1), (3) potentiation of recurrent synaptic strengths from excitatory neurons to *PV*^+^-interneurons *(ζ_PE_* > 1), and (4) potentiation of recurrent synaptic strengths from *PV*^+^ to excitatory neurons (*ζ_EP_* > 1).

**Table 1:** Default parameters of the single neurons and networks, and parameters for modeling the synaptic changes during monocular deprivation in layer 4 of V1m (Figs. 1–4) as well as for the changes during whisker deprivation in layer 2/3 of S1 (Fig. 6). EPSG/IPSG: excitatory/inhibitory postsynaptic conductance; LGN: lateral geniculate nucleus; BKG: background.

| Symbol | Value/<br>Range | Description |
| --- | --- | --- |
| $C_m$ | 200 pF | Membrane capacitance |
| $g_L$ | 10 nS | Leak conductance |
| $E_L$ | -70 mV | Leak reversal potential |
| $V_\theta$ | -50 mV | Threshold potential |
| $V_r$ | -58 mV | Post-spike reset potential |
| $E_E$ | 0 mV | Excitatory reversal potential |
| $E_P, E_S$ | -85 mV | Inhibitory reversal potential from $PV^+$ ($P$) and $SST^+$ ($S$) interneurons |
| $\tau_{syn}$ | 5 ms | Synaptic conductance time constant |
| $N_E$ | 4000 | Number of excitatory neurons |
| $N_P$ | 1000 | Number of $PV^+$-interneurons |
| $N_S$ | 500 | Number of $SST^+$-interneurons |
| $p$ | 0.1 | Connection probability between two cells |
| $J$ | 0.1 nS | EPSP amplitude $E \rightarrow E$, $E \rightarrow PV^+$ and $E \rightarrow SST^+$ |
| $g_{rc}J$ | 0.8 nS | IPSP amplitude $PV^+ \rightarrow E$ and $PV^+ \rightarrow PV^+$ |
| $K$ | 1.6 nS | IPSP amplitude for $SST^+ \rightarrow E$ and $SST^+ \rightarrow PV^+$ connections |
| $J_s, J_b$ | 0.5 nS | EPSP amplitude $LGN, BKG \rightarrow E$, $BKG \rightarrow SST^+$ and scale for EPSP $LGN \rightarrow PV^+$ |
| $g_{fw}$ | 2 | Multiplicative factor for strong feedforward EPSP $LGN \rightarrow PV^+$ |
| $R_{stim}$ | 1000 Hz | Rate of feedforward input spike trains $LGN \rightarrow E$ and $LGN \rightarrow PV^+$ and of background input spike trains $BKG \rightarrow E$ and $BKG \rightarrow SST^+$ |
| $\delta_E$ | [0.5, 1.0] | Depression of feedforward synapses onto excitatory neurons |
| $\delta_P$ | [0.5, 1.0] | Depression of feedforward synapses onto $PV^+$-interneurons |
| $\zeta_{PE}$ | [1.0, 1.5] | Potentiation of recurrent synapses from excitatory neurons to $PV^+$-interneurons |
| $\zeta_{EP}$ | [1.0, 1.5] | Potentiation of recurrent synapses from $PV^+$-interneurons to excitatory neurons |
| $\rho_{EI}$ | [1.0, 1.5] | E/I ratio of feedforward synaptic changes (increased in V1) |
| $\xi_\theta$ | [0, 3] mV | Firing threshold of $PV^+$-interneurons (increased in S1) |

We investigated the effect of these feedforward and recurrent synaptic changes on excitatory and inhibitory activity separately. For each pair of feedforward (*δ_E_,δ_P_*) or recurrent (*ζ_EP_,ζ_PE_*) synaptic changes, we generated a plane of firing rate changes in the excitatory and *PV*^+^ populations relative to their respective baseline firing rates before plasticity (Fig. 1B, C). We found that depression of feedforward synaptic strengths, which reduces the feedforward drive to the network, can have either facilitating or suppressive effects on the activity of excitatory neurons and *PV*^+^-interneurons (Fig. 1B). A linear relationship between *δ_E_* and *δ_P_* separates the facilitation and suppression areas of excitatory and *PV*^+^ firing rates. The slope of this line is determined by the relatively stronger feedforward drive onto *PV*^+^-interneurons compared to excitatory neurons (*g_fw_*, Fig. 1A and Methods), determined experimentally (Ji et al., 2016; Pfeffer et al., 2013). To capture the relationship between the relative excitatory and inhibitory feedforward drive, we introduced the E/I ratio of feedforward synaptic changes, *ρ_EI_* = *δ_E_/δ_P_*. This ratio has been measured to be bigger than unity after brief MD from the thalamus to layer 4 (Miska et al., 2018), and also from layer 4 to layer 2/3 in the visual cortex (Kuhlman et al., 2013). Therefore, we found that synaptic changes exclusively in the feedforward pathway from the thalamus to cortex, which increase the E/I-ratio, increase both excitatory and inhibitory firing rates (Fig. 1B). In contrast, purely recurrent synaptic changes measured in layer 4 of V1 after two days of MD (Maffei et al., 2006; Miska et al., 2018), which potentiate synaptic strengths from excitatory neurons to *PV*^+^-interneurons (*ζ_PE_* > 1), and from *PV*^+^-interneurons to excitatory neurons (*ζ_EP_* > 1), have a purely suppressive effect on the excitatory and inhibitory firing rates (Fig. 1C). Combining feedforward and recurrent plasticity can either increase or decrease excitatory and inhibitory firing rates, depending on whether feedforward or recurrent synaptic changes dominate (Fig. 1D).

Intriguingly, modeling feedforward and recurrent plasticity separately reveals that firing rate changes of excitatory neurons and *PV*^+^-interneurons should tightly follow each other in the entire parameter space of feedforward and recurrent synaptic changes (Fig. 1B-D). This implies that if inhibitory firing rates decrease at MD1, so should excitatory firing rates (when in fact, experimentally, they stay at baseline), and if excitatory firing rates decrease at MD2, so should inhibitory firing rates (when in fact, experimentally, they recover back to baseline). Hence, this result is inconsistent with the independent modulation of excitatory and *PV*^+^ firing rates following brief MD observed *in vivo* (Hengen et al., 2013), suggesting that other factors might be at play.

### Network response to MD-induced synaptic changes depends on coupling scale

We next sought to identify a plausible mechanism behind the lack of independent modulation of firing rate changes in the excitatory and *PV*^+^ populations. Of the two parameters which determine recurrent synaptic strengths in the model network, the overall coupling scale, *J*, is more important than the relative scale of the inhibitory synaptic strength, *g_rc_* in determining the network response to MD-induced plasticity (Table 1). The parameter *J* has been extensively studied as a determinant of the operating regime of the network (Brunel, 2000; Hennequin et al., 2018; Murphy and Miller, 2009; Renart et al., 2010; van Vreeswijk and Sompolinsky, 1996). A network with a strong coupling scale *J* operates in a regime where the excitatory dynamics are stabilized by recurrent inhibition *(inhibition stabilized network,* ISN). In contrast, a network with weak *J* operates in a regime where the excitatory dynamics are stable without recurrent inhibition (non-ISN) (Tsodyks et al., 1997). We found two very different behaviors in the network at the opposite ends of the coupling scale, *J*. While at high coupling, excitatory and *PV*^+^ responses show a tight alignment (Fig. 1 with *J* = 0.1 nS), for weak to intermediate coupling when the network is in the non-ISN regime, excitatory and *PV*^+^ firing rates can be independently modulated for all feedforward and recurrent synaptic changes (Supplementary Fig. S1 with *J* = 0.01 nS). Hence, the transition from independent to tight modulation of excitatory and inhibitory firing rates corresponds to the transition from non-ISN to ISN with increasing coupling scale *J*.

To quantify how closely the firing rates of excitatory and inhibitory neurons follow each other as a function of *J*, we computed the fractional response area of facilitation for excitatory neurons *(F_E_*) and for *PV*^+^-interneurons *(F_P_*), and the overlap between excitatory and *PV*^+^ response areas *(O_EP_*, Fig. 2A). We did this for each parameter space of feedforward (*δ_E_, δ_P_*) (Fig. 2B), recurrent *(ζ_EP_, ζ_PE_*) (Fig. 2C), and combined *(ζ_PE_,ρ_EI_*) (Fig. 2D) synaptic changes. In all cases, we found that the facilitation area for *PV*^+^-interneurons, and consequently the overlap between excitatory and *PV*^+^ response areas, abruptly changes at a critical value of coupling (*J* ≈ 0.017 nS). For instance, in the parameter space of feedforward changes, *PV*^+^ facilitation emerges at *J* ≈ 0.017 nS (*F_P_*, Fig. 2B), while in the parameter space of recurrent changes, *PV*^+^ facilitation decreases to 0 at the same critical *J* (*F_P_*, Fig. 2C).

**Figure 2:**
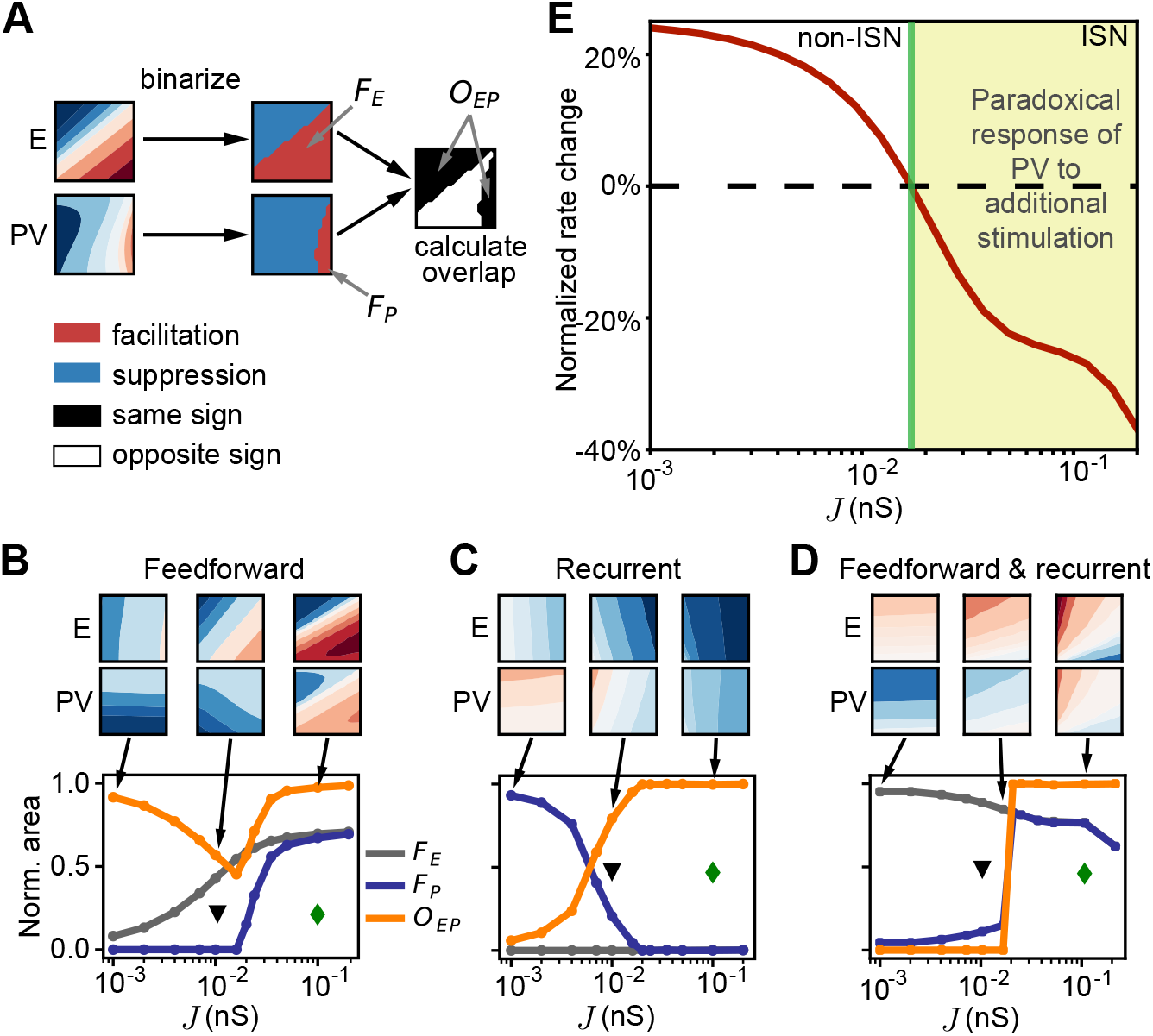
Firing rate changes in response to MD depend on the network coupling scale J. **(A)** Schematic to determine the overlap of facilitating and suppressive response areas of excitatory neurons and *PV*^+^-interneurons. The thresholded plane of responses (Fig. 1) for both neuron types is used to compute the total facilitating area and quantify how closely excitatory and *PV*^+^ firing rates follow each other through the overlap of facilitation and suppression. **(B)** Network with feedforward depression only (Fig. 1B). Fractional area of facilitation for excitatory neurons (*F_E_*, gray), *PV*^+^-interneurons (*F_P_*, blue) and the overlap between excitatory and *PV*^+^ response areas (*O_EP_*, orange) as a function of the overall coupling scale J. Green diamond shows the value of *J* used in Fig. 1 (ISN), black triangle shows the value of *J* used in Supplementary Fig. S1 (non-ISN). **(C)** Same as (B) for recurrent potentiation only (Fig. 1C). **(D)** Same as (B) for combined feedforward and recurrent plasticity (Fig. 1D). **(E)** Normalized rate change of *PV*^+^-interneurons in response to additional input to them as a function of coupling scale J. Additional input is implemented as an increase in *δ_P_* (from 1 to 1.1).

Using a reduced linear population rate model (Supplementary Fig. S2) (Tsodyks et al., 1997; Wilson and Cowan, 1972), we found that a key network property related to the transition from non-ISN to ISN is the emergence of facilitation in *PV*^+^-interneurons at the critical coupling *J* following feedforward plasticity (Methods). This transition is a direct consequence of the emergence of the *paradoxical effect* at the critical *J*, where in response to additional excitatory drive, the inhibitory population paradoxically decreases its rate together with the excitatory population (Fig. 2E) (Tsodyks et al., 1997). Therefore, the emergence of the paradoxical effect in the model network with excitatory and a single type of inhibitory neurons can explain the transition from independent to tight modulation of excitatory and inhibitory firing rates as a function of the coupling scale.

These results argue that only the non-ISN supports the experimentally observed independent modulation of excitatory and inhibitory firing rates in response to synaptic changes after brief MD due to the absence of the paradoxical effect. However, cortical circuits seem to operate in the ISN regime (Lien and Scanziani, 2013; Ozeki et al., 2009; Sanzeni et al., 2020), which due to the paradoxical effect cannot explain the independent modulation of excitatory and inhibitory firing rates after brief MD. Moreover, in the non-ISN, the magnitude of this modulation to this plasticity is much weaker, suggesting that experimentally measurable changes in firing rates might require synaptic changes larger than what seems biologically plausible.

### Network model with two subtypes of interneurons

Given the role of interneuron diversity in a range of cortical functions, including disinhibition, response reversal, and surround suppression, we next added the second-largest class of cortical interneurons, *SST*^+^-interneurons, to the ISN (Tremblay et al., 2016; Urban-Ciecko and Barth, 2016). We modeled *SST*^+^-interneurons which project to excitatory neurons and *PV*^+^-interneurons via the coupling parameter *K*, but which do not receive inhibition from *PV*^+^-interneurons nor inhibit each other (Fig. 3A) (Pfeffer et al., 2013; Xu et al., 2013). *SST*^+^-interneurons receive almost no thalamic feedforward input (Ji et al., 2016), but receive feedback from higher-order cortical centers and integrate inputs over large areas in the visual cortex (Adesnik et al., 2012; Batista-Brito et al., 2018; Tremblay et al., 2016; Zhang et al., 2014). We modeled these inputs from outside the local patch of cortex represented by the network as a background that provides input to both *SST*^+^ and excitatory neurons (Fig. 3A). Upon introducing *SST*^+^-interneurons, the ISN remained in an asynchronous and irregular firing state independent of the *SST*^+^ feedback *K* (Fig. 3B). We found that the overall effect of *K* on the average rate is suppressive (Fig. 3C), although *SST*^+^ feedback is symmetric providing direct inhibition to excitatory neurons as well as indirect disinhibition to excitatory neurons via *PV*^+^-interneurons. We next studied whether the ISN with these two subtypes of inhibitory interneurons is a suitable candidate to explain the independent modulation of excitatory and inhibitory firing rates.

**Figure 3:**
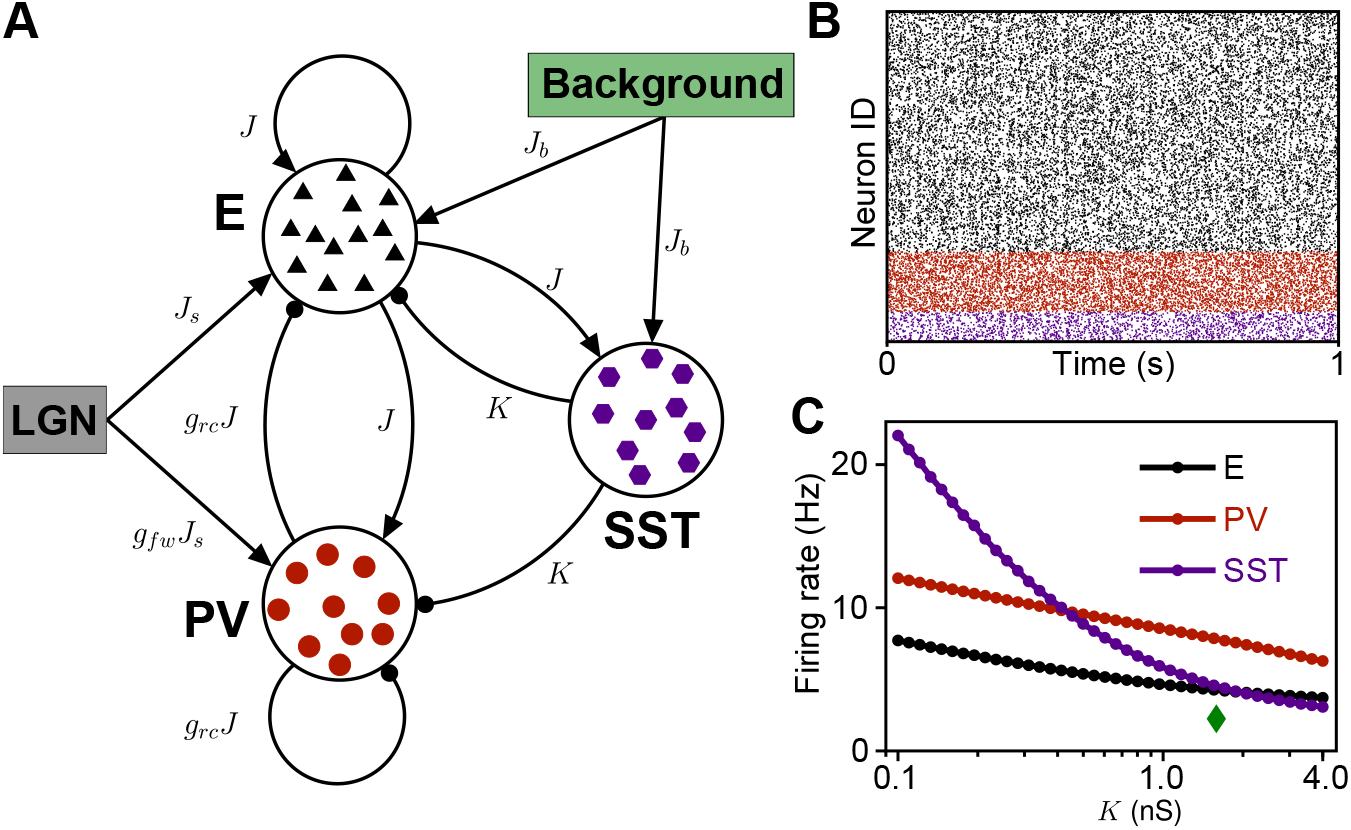
Network structure and spiking activity in the model with two subtypes of interneurons. **(A)** Schematic of the connectivity in the network with two subtypes of interneurons. **(B)** Spike raster of activity in the network with *K* = 1.6 nS. **(C)** Firing rates as a function of feedback strength *K*. The green diamond shows the value of K used in (B).

### *SST*^+^ -interneuron feedback can invert *PV*^+^-interneuron responses

Implementing feedforward synaptic changes in this ISN with two inhibitory subtypes, we found that the excitatory response is qualitatively similar compared to the network without *SST*^+^ feedback (Fig. 4A vs. Fig. 1B). Strikingly, the presence of strong *SST*^+^ feedback *K* inverts the response of *PV*^+^-interneurons in the (*δ_E_, δ_P_*) plane of synaptic changes (Fig. 4A vs. Fig. 1B). As before, to quantify the joint modulation of excitatory and *PV*^+^ firing rates, we computed the fractional response area of facilitation for excitatory neurons (*F_E_*) and for *PV*^+^-interneurons (*F_P_*), and the overlap between excitatory and *PV*^+^ response areas (*O_EP_*), now as a function of *K* (Fig. 4D). We observed that the inversion of *PV*^+^ responses occurs for *SST*^+^ feedback bigger than a critical value (*K* ≈ 0.6 nS), where the facilitation area of *PV*^+^ responses achieves a minimum as a function of *K* (*F_P_*, Fig. 4D).

**Figure 4:**
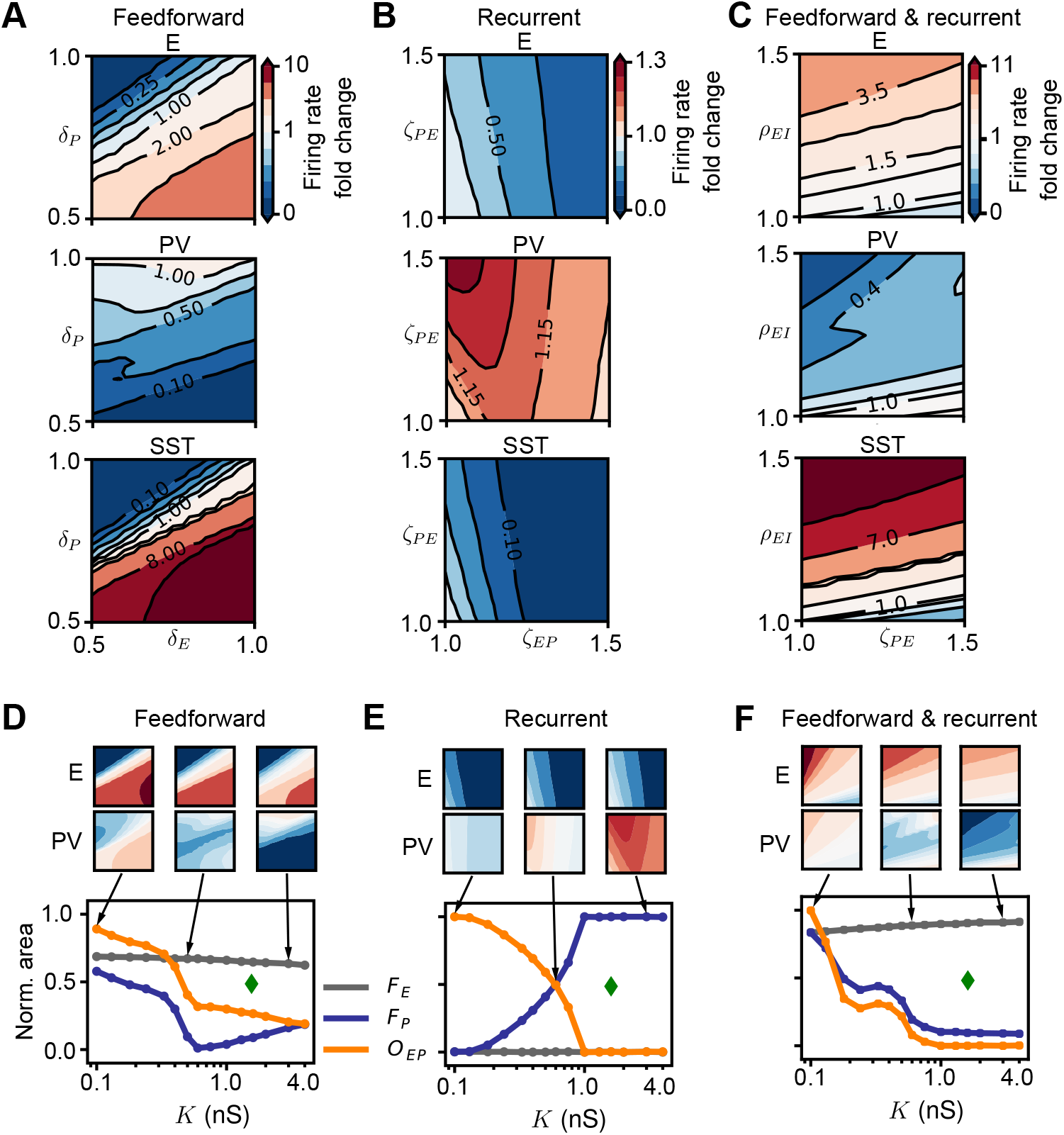
*SST*^+^ feedback selectively inverts *PV*^+^ responses. **(A)** Network firing rate in the (*δ_E_, δ_P_*) plane as fold-change of baseline firing rate (baseline in top right corner) for excitatory (top panel), *PV*^+^ (middle panel) and *SST*^+^neurons (bottom panel) in the network with strong *SST*^+^ feedback (*K* = 1.6 nS). **(B)** Network firing rate in the *(ζ_EP_, ζ_PE_*) plane as fold-change of baseline firing rate (baseline in bottom left corner). **(C)** Network firing rate in the *(ζ_PE_, ρ_EI_*) plane as fold-change of baseline firing rate (baseline in bottom left corner). **(D)** Network with feedforward depression only. Fractional area of facilitation for excitatory neurons (*F_E_*, gray), *PV*^+^-interneurons (*F_P_*, blue) and the overlap between excitatory and *PV*^+^ response areas (*O_EP_*, orange). **(E)** Same as (D) for recurrent potentiation only. **(F)** Same as (D) for combined feedforward and recurrent plasticity. The green diamond in (D)-(F) shows the value of K used in (A)-(C).

The inversion of *PV*^+^ responses in this ISN with two inhibitory subtypes can be explained by a generalization of the paradoxical effect. Using the reduced linear population rate model (Supplementary Fig. S3), we can explicitly derive the link between the inversion of *PV*^+^-responses to feedforward synaptic changes and the reversal of the paradoxical effect as a function of the *SST*^+^ feedback *κ* in the reduced model (Eq. 30). As *SST*^+^ feedback increases, *PV*^+^-interneurons start to exhibit non-paradoxical facilitation rather than paradoxical suppression in response to a step-current (Fig. 5A). For any value of *SST*^+^ feedback, *κ*, the *PV*^+^ population activity initially transiently increases, leading to a decrease of excitatory population activity (Fig. 5B). For small *κ*, the decrease of excitatory activity suppresses the steady-state *PV*^+^ activity below baseline, leading to the classical paradoxical effect (Fig. 5B, left). As the *SST*^+^ feedback, *κ*, increases, the steady-state *PV*^+^ activity is not only dictated by the dwindling of recurrent excitation as in the case of the paradoxical effect. Rather, *PV*^+^ activity becomes dominated by the release from inhibition mediated through *SST*^+^. Hence, the paradoxical decrease of *PV*^+^ activity is only transient, followed by a recovery of *PV*^+^ activity back to baseline at an intermediate value of *κ* (Fig. 5B, middle). For large *κ*, the steady-state *PV*^+^ activity increases above baseline (Fig. 5B, right).

**Figure 5:**
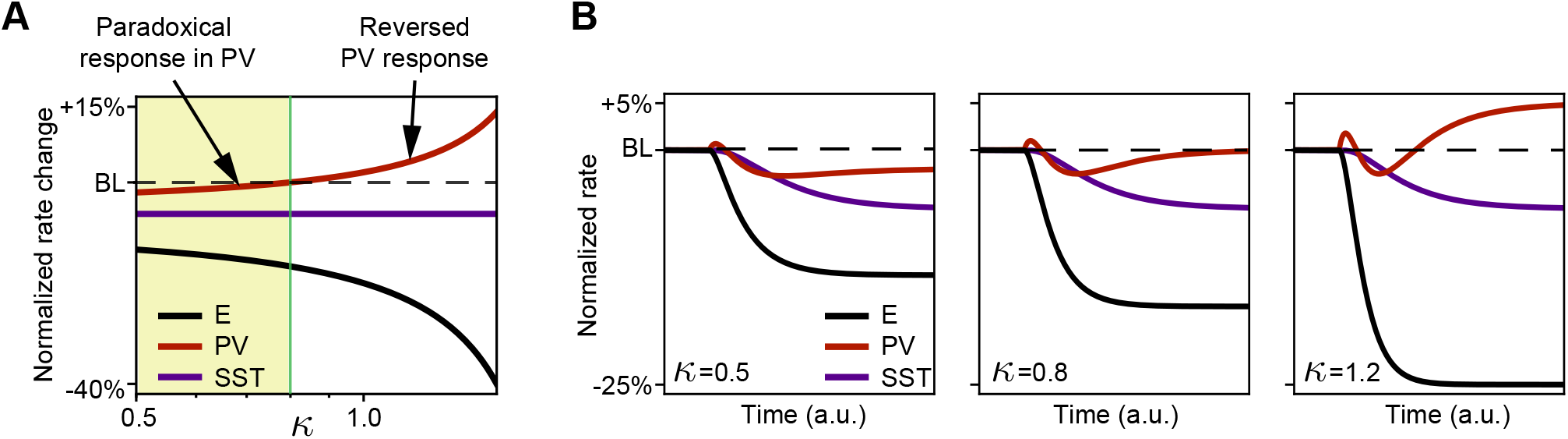
The inversion of responses of *PV*^+^-interneurons can be explained by reversal of the paradoxical effect. **(A)** Relative change of steady-state rate induced by current injection to *PV*^+^-interneurons as a function of *SST*^+^ feedback *κ*. The rate of each population is normalized to its baseline firing rate before additional current injection (BL). **(B)** Dynamics of the linear population rate model following onset of step current to *PV*^+^-interneurons for *κ* = 0.5 (left), *κ* = 0.8 (middle) and *κ* = 1.2 (right). Colors are the same as in (A).

Recurrent plasticity continues to have a strong suppressive effect on the excitatory firing rates, accompanied by a similar response in the *SST*^+^-interneurons (Fig. 4B vs. Fig. 1C). As the paradoxical effect reverses, *PV*^+^-interneurons invert their response, exhibiting pure facilitation for large values of *K* (Fig. 4B vs. Fig. 1C). As a result of this inversion of *PV*^+^ responses, the overlap between the excitatory and *PV*^+^ responses decays to zero (*O_EP_*, Fig. 4E). Combining feedforward and recurrent synaptic changes further corroborates the inversion of *PV*^+^ responses with increasing *SST*^+^ feedback (Fig. 4C, F).

We conclude that, in strongly coupled networks operating in the ISN regime typical of the sensory cortex, sufficiently strong feedback from *SST*^+^-interneurons can invert the responses of *PV*^+^-interneurons following feedforward and recurrent synaptic changes observed in early MD. This inversion of *PV*^+^ responses provides a natural substrate for the independent modulation of excitatory and inhibitory firing rates as observed *in vivo* during brief MD (Hengen et al., 2013).

### Sensory perturbation induces similar activity changes in somatosensory cortex via different plasticity mechanisms

Similar to MD, whisker deprivation (WD) is a sensory deprivation, where plucking a subset of the whiskers on one cheek of the animal affects the barrel cortex of the rodent primary somatosensory cortex (S1). Interestingly, brief WD induces different changes in the circuitry of S1 than brief MD does in V1 (Gainey et al., 2018; House et al., 2011; Li et al., 2014). Rather than feedforward synaptic depression, a reduction of feedforward inhibition emerges from a decrease in the intrinsic excitability of *PV*^+^-interneurons through an increase of their firing threshold (Gainey et al., 2018). This leads to a similar increase in the feedforward excitation to inhibition ratio as in V1 (Kuhlman et al., 2013; Li et al., 2014; Miska et al., 2018). Our modeling approach allowed us to investigate whether these different types of plasticity in V1 and S1 during sensory deprivation change the modulation of excitatory and inhibitory firing rates in a corresponding microcircuit model of S1 (Fig. 6A).

**Figure 6:**
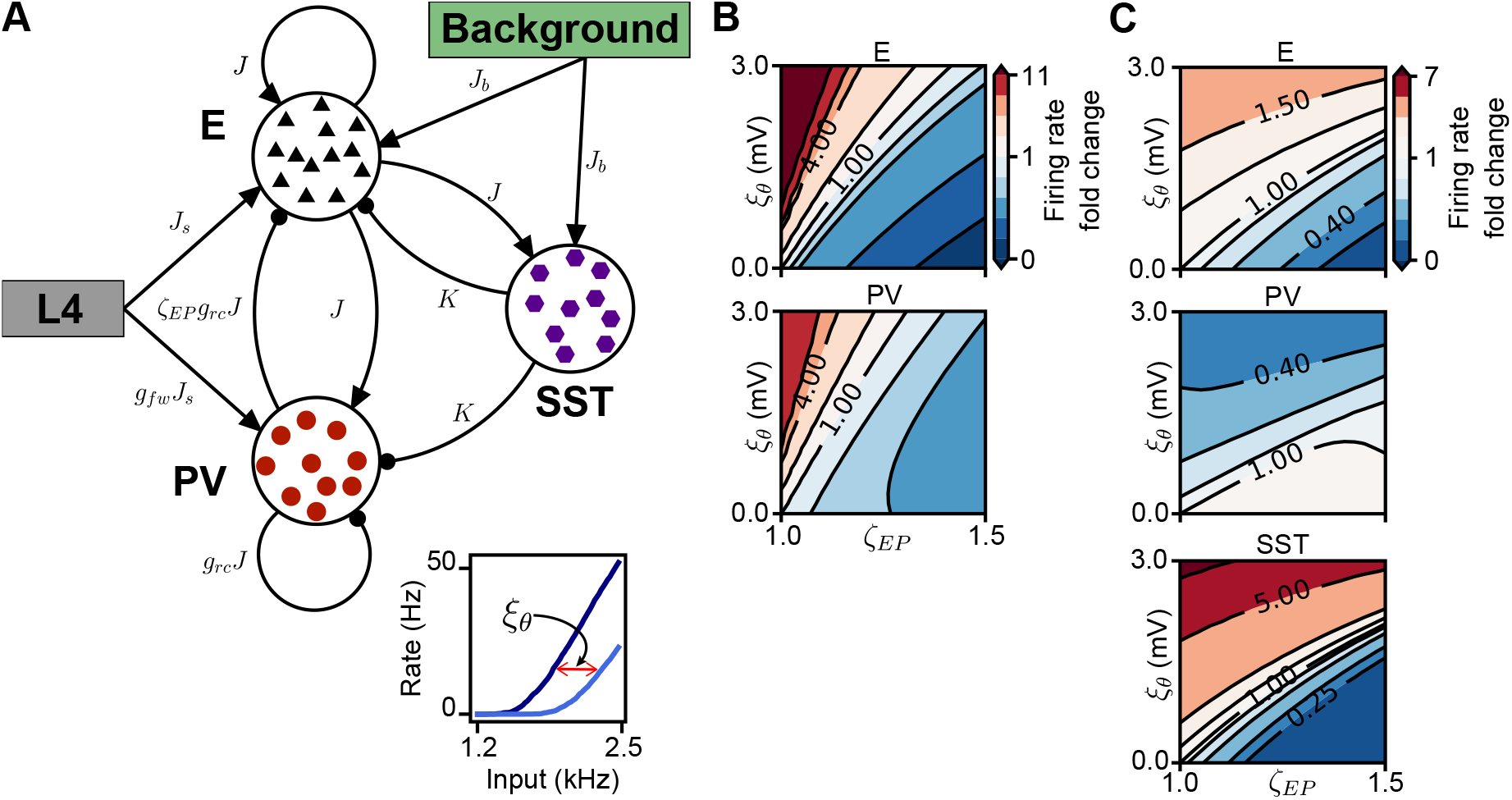
Combining feedforward and recurrent plasticity in S1. **(A)** Schematic for combined recurrent and feedforward plasticity in S1: shift in recurrent E/I-ratio through *ζ_EP_*, shift in feedforward E/I-ratio through increasing firing threshold of *PV*^+^-interneurons (*ξ_θ_*). Inset: The f-I curve of a single leaky integrate-and-fire neuron for baseline firing threshold (dark blue) and with increased firing threshold (light blue). The excitability of *PV*^+^-interneurons is decreased via the parameter *ξ_θ_*. **(B)** Network firing rate as fold-change of baseline firing rate (baseline in bottom left corner) in *(ζ_EP_, ξ_θ_*) plane in the network with only one type of interneurons (*PV*^+^). **(C)** Network firing rate as fold-change of baseline firing rate in *(ζ_EP_, ξ_θ_*) plane in the network with two subtypes of interneurons (*PV*^+^ and *SST*^+^), with strong *SST*^+^ feedback *K* = 1.6 nS.

As before, we first considered a network with a single interneuron type by setting the *SST*^+^ feedback, *K*, to zero. We modeled the decrease of the intrinsic excitability of *PV*^+^-interneurons by increasing their firing threshold by *ξ_θ_* (Fig. 6A, inset). In an ISN model of S1 with only *PV*^+^-interneurons, increasing the firing threshold of *PV*^+^-interneurons facilitates their firing rates, just like the depression of the feedforward synaptic strength from the LGN onto excitatory and inhibitory neurons in the model V1 network. This decrease of intrinsic excitability of *PV*^+^-interneurons in S1 was proposed as a home-ostatic mechanism that counteracts the lack of stimulation after deprivation and enhances firing rates (see also House et al. (2011); Li et al. (2014)).

We further implemented potentiation of recurrent inhibition as in V1 by *ζ_PE_*. Experimentally, brief (1 day of) WD in S1 has been found to exhibit a tendency but no statistical significance for the potentiation of recurrent inhibitory synapses from *PV*^+^ to excitatory neurons (Gainey et al., 2018). In contrast, prolonged (6-12 days of) WD has shown a pronounced increase in this connection strength (House et al., 2011). Hence, the potentiation of recurrent inhibition by *ζ_PE_* seems to counteract the facilitation of activity by *ξ_θ_*, giving rise to the same antagonistic regulation we found for early MD-induced plasticity in V1 (Fig. 6B, compare to Fig. 1D). Similar to MD-induced plasticity, excitatory and *PV*^+^ activity changes in this network with WD-induced plasticity are tightly coordinated in the ISN. Importantly, after introducing strong feedback from *SST*^+^-interneurons, the response of *PV*^+^-interneurons again inverts relative to the excitatory responses due to the reversal of the paradoxical effect (Fig. 6C).

Therefore, our modeling demonstrates that the dissimilar circuit changes induced by sensory deprivation (WD or MD) in two different sensory cortices, which decrease intrinsic excitability of *PV*^+^-interneurons or depress feedforward drive to *PV*^+^-interneurons, respectively, lead to similar regulation of overall activity as they interact with potentiation of recurrent inhibition. Hence, excitatory and inhibitory firing rates in S1 during brief WD could be independently modulated similar to those measured in V1 (Hengen et al., 2013) in the presence of feedback from a second inhibitory subtype due to the reversal of the paradoxical effect in an ISN.

## Discussion

We investigated how synaptic changes induced by sensory deprivation affect the firing rates of excitatory and inhibitory neurons in a microcircuit network model of the sensory cortex. Specifically, we modeled two pathways: (1) the reduction of feedforward excitability manifested as a depression of feedforward, thalamocortical synapses onto excitatory and inhibitory neurons observed following monocular deprivation in visual cortex (or as a decrease of intrinsic excitability in inhibitory neurons following whisker deprivation in somatosensory cortex), and (2) the potentiation of recurrent synapses between excitatory and inhibitory neurons. Crucially, we found that strong feedback from a second interneuron subtype, *SST*^+^-in addition to *PV*^+^-interneurons, is needed to explain the independent modulation of excitatory and inhibitory activity observed experimentally in the primary visual cortex during the critical period of plasticity.

Previous studies have investigated MD-induced plasticity in V1, but with a focus on ocular dominance plasticity in binocular cortex governed by winner-take-all competition where the open eye out-competes the closed eye (Blais et al., 2009, 1999; Kuhlman et al., 2013; Miller et al., 1989; Toyoizumi et al., 2014; Toyoizumi and Miller, 2009; Toyoizumi et al., 2013). Since most brain regions do not exhibit such strong competition, we focused on plasticity in the monocular region of primary visual cortex where synaptic and activity changes during MD have recently been measured (Hengen et al., 2013, 2016; Miska et al., 2018). Some models have already begun to incorporate these experiments by implementing a careful orchestration of biologically inspired synaptic and homeostatic plasticity rules to explain the recovery of diverse aspects of network dynamics after MD (Ma et al., 2019; Sweeney et al., 2018; Wu et al., 2020). Although our approach is agnostic about the plasticity rules underlying deprivation-induced synaptic changes, it offers the advantage to turn individual synaptic changes ‘on’ or ‘off’ in different network regimes. This gives us complete control to study the effect of the timing of individual plasticity mechanisms in the network.

We found that the operating regime of the network, determined by the overall coupling scale, has a major impact on how deprivation-induced plasticity shapes firing rate changes. Inhibition stabilized networks (ISNs) with a single inhibitory subtype and strong coupling, commonly used to describe the cortex (Mahrach et al., 2020; Sanzeni et al., 2020), cannot capture the independent modulation of excitatory and inhibitory firing rates after brief MD due to the presence of the paradoxical effect. We propose that sufficiently strong feedback from *SST*^+^-interneurons can achieve the independent modulation of excitatory and inhibitory firing rates in response to MD-induced synaptic changes by reversing the paradoxical effect. This is consistent with previous theoretical results for a network with multiple subtypes of interneurons, where a specific inhibitory population can behave non-paradoxically in response to an input current, even if the whole network operates as an ISN (Garcia del Molino et al., 2017; Litwin-Kumar et al., 2016).

An alternative to changing the overall coupling scale is to change the response gain of single neurons in the network, as is the case in the stabilized supralinear network (SSN). The SSN has recently gained attention as a circuit model of V1 with powerful computational capabilities, including contrast gain control and nonlinear response amplification (Ahmadian et al., 2013; Rubin et al., 2015). Our systematic analysis over a wide range of coupling scales could be interpreted as the local linear approximation of the SSN in response to MD-induced synaptic changes. Hence we expect all results to hold also in the SSN.

Our microcircuit model with two interneuron types suggests that *PV*^+^-interneurons in layer 4 in monocular V1 should respond non-paradoxically upon optogenetic stimulation. However, recent experiments find that *PV*^+^-interneurons in layer 2/3 and upper layer 4, as well as deep layers of V1, show a paradoxical response following stimulation (Sanzeni et al., 2020). It is possible that our proposed non-paradoxical response of *PV*^+^-interneurons is a transient property of the developing cortex during the critical period that disappears with maturation, e.g. through a developmental decrease of initially strong coupling from *SST*^+^-onto *PV*^+^-interneurons (Tuncdemir et al., 2016), or that it depends on the anatomical region or the cortical layer (Adesnik, 2017; Atallah et al., 2012; Kato et al., 2017; Mahrach et al., 2020; Sadeh et al., 2017).

Our results make concrete predictions that can be tested with further experiments that characterize the population activity in response to driving individual circuit elements at different stages during development and in different layers. First, we predict that *SST*^+^-interneurons in layer 4 during the critical period should decrease their activity in response to additional stimulation of *PV*^+^-interneurons (e.g. through optogenetic activation). This decrease would be the result of a withdrawal of recurrent excitation, rather than direct inhibition from *PV*^+^-to *SST*^+^-interneurons, and could specifically be investigated by experimentally measuring the respective conductances in individual *SST*^+^-interneurons. A second, related prediction is that the firing rates of *SST*^+^-interneurons during the first two days of MD should be tightly coordinated with excitatory firing rates. Separating excitatory and *SST*^+^ firing rates is currently difficult in *in vivo* extracellular recordings because, depending on the morphological subtype of the *SST*^+^-interneurons, the spike-waveform would either be classified in the regular-spiking or the fast-spiking type, but chronic calcium imaging recordings present a possible alternative (Rose et al., 2016). A further unknown is the plasticity of recurrent connections involving *SST*^+^-interneurons in L4 during MD (Scheyltjens and Arckens, 2016; Urban-Ciecko and Barth, 2016). Although the plasticity of synapses involving electrophysiologically identified regular spiking non-pyramidal units has been measured (Maffei et al., 2004), these cells were not genetically labeled as *SST*^+^, and the measurements were performed before the onset of the critical period, which might involve a different set of plasticity mechanisms than the critical period (Lefort et al., 2013). Third, we modeled the potentiation of intracortical inhibition during brief MD as a change in the excitatory drive from excitatory neurons to *PV*^+^-interneurons to be consistent with known plasticity mechanisms (Maffei et al., 2006; Miska et al., 2018). However, it is possible that the additional sources of inhibition from *SST*^+^-interneurons contribute to this increase in inhibition. For instance, potentiation of synaptic strength from *SST*^+^-onto excitatory neurons and/or depression of synaptic strength from *SST*^+^-onto *PV*^+^-interneurons could also underlie the increase in intracortical inhibition. We found that implementing these changes in our recurrent network generates comparable results to the firing rate modulation of excitatory and *PV*^+^-interneurons (Fig. S4), suggesting the existence of a parallel, possibly redundant pathway to generate the same firing rate changes.

In summary, our mechanistic modeling of a network with more than one interneuron type suggests the following sequence as the most parsimonious explanation for the independent modulation of excitatory and inhibitory firing rates after sensory deprivation observed experimentally (Fig. 7). First, during the first day of deprivation, depression of the feedforward synaptic weights from LGN to excitatory neurons and *PV*^+^-interneurons in the visual cortex (or of *PV*^+^ excitability in the somatosensory cortex), can give rise to strong suppression of *PV*^+^ firing rates, while excitatory rates remain at baseline, only in the ISN with strong *SST*^+^-feedback (Fig. 7, left). Instead, in the ISN without *SST*^+^-feedback the firing rates of both excitatory and *PV*^+^ populations facilitate or suppress together. Second, during the second day of deprivation, potentiation of recurrent inhibition in the cortex can strongly suppress excitatory firing rates and recover *PV*^+^ firing rates to baseline through potentiation of activity from the earlier suppressed state (Fig. 7, right). Again, this is only possible in the ISN with strong *SST*^+^-feedback which reverses the paradoxical effect in *PV*^+^-interneurons, while in the ISN without *SST*^+^-feedback the presence of the paradoxical effect leads to a tight coordination of excitatory and *PV*^+^ firing rates. More generally, our work provides a critical link between changes on the single-synapse level and activity modulation on the circuit level during perturbation of normal development as a function of cortical operating regime and interneuron diversity.

**Figure 7:**
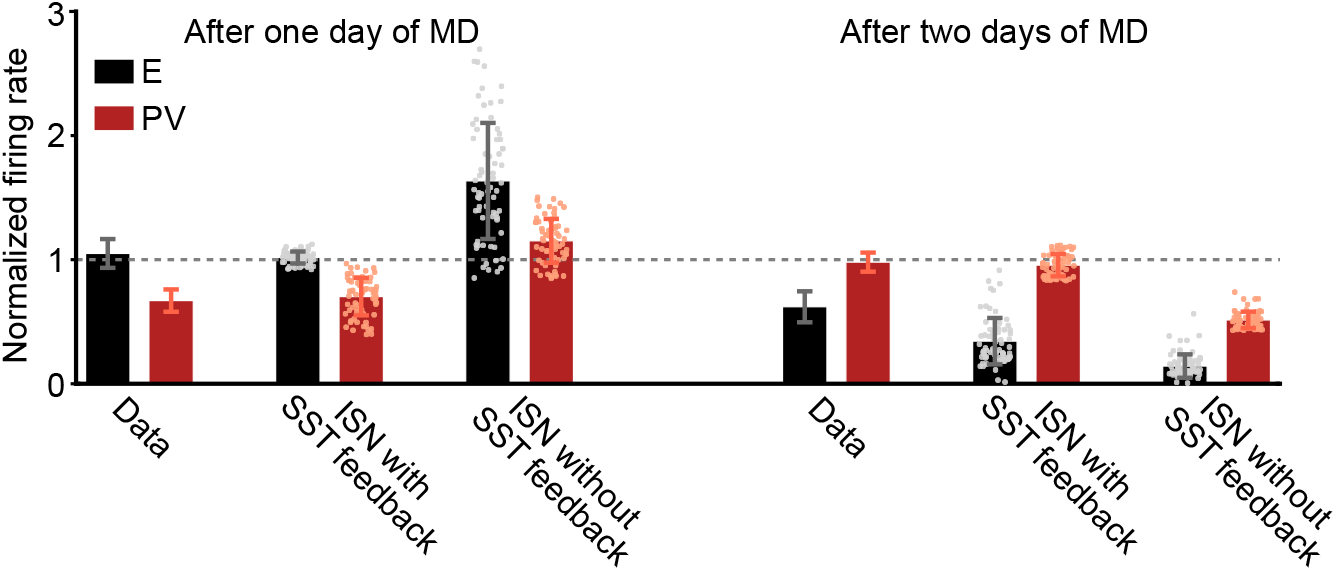
An ISN with strong *SST*^+^-feedback can capture the independent modulation of excitatory and inhibitory firing rates observed during brief MD. Firing rates of excitatory (black) and *PV*^+^ (red) neurons after one day of MD (left histograms) and after two days of MD (right histograms), normalized by the firing rates before MD-induction. Data are extracted from (Hengen et al., 2013) and reproduced in the spiking network in the ISN regime that includes strong feedback from *SST*^+^ and in an ISN without *SST*^+^-feedback (Methods). Experimental data are mean ± SEM, simulation data are mean ± SD.

## Acknowledgments

We thank all members of the Gjorgjieva lab for discussions, Gina Turrigiano for fruitful discussions throughout the project and critical feedback on the manuscript, and Matthias Kaschube and Hiroshi Ito for additional comments on the manuscript. This work was supported by the Max Planck Society (LMAR, JG). This project has received funding from the European Research Council (ERC) under the European Union’s Horizon 2020 research and innovation program (Grant agreement No. 804824).

## Author Contributions

LMAR and JG designed the research and developed the model. LMAR analyzed the model and performed simulations. LMAR and JG prepared the figures and wrote the manuscript.

## Data Availabilty

The code used for model simulations is available at GitHub (https://github.com/comp-neurai-circuits/MD-dynamics).

## Methods

### Spiking network model

We studied recurrently connected networks of leaky integrate-and-fire neurons (LIF), consisting of *N_E_* excitatory neurons, *N_P_ PV*^+^-interneurons and *N_S_ SST*^+^-interneurons. The membrane potential *V*(*t*) of a single neuron follows the dynamics:

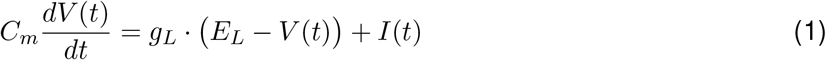

where *C_m_* is the membrane capacitance, *g_L_* the conductance of the leak-current, *E_L_* the leak reversal potential and *I*(*t*) the total input current (parameter values provided in Table 1). When the membrane potential reaches a firing threshold *V_θ_*, a spike is emitted and the potential is reset to *V_r_*. The neurons are randomly and sparsely connected with connection probability p with fixed in-degree by conductance-based synapses, or weights. Thus, all values for synaptic weights are positive and the excitatory or inhibitory action of a presynaptic neuron is determined by the respective reversal potential (Table 1). All excitatory weights are denoted by *J* regardless of the postsynaptic target (Brunel, 2000); this parameter is known as the coupling scale and determines the operating regime of the network (Tsodyks et al., 1997). Synapses from *PV*^+^-interneurons are scaled by the factor *g_rc_* (Brunel, 2000). Synapses from *SST*^+^-interneurons are denoted by *K*. Hence, the population connectivity matrix is given by:

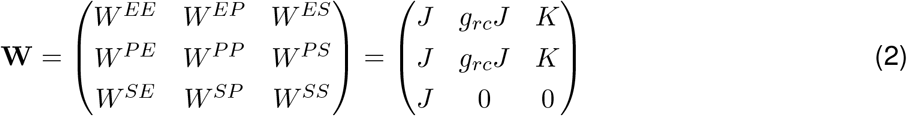

where *W^AB^* denotes the connection from population *B* to population *A*, and *E* denotes excitatory neurons, *P* denotes *PV*^+^-interneurons and *S* denotes *SST*^+^-interneurons. The case with only two populations, excitatory and *PV*^+^, can be derived from this by setting the *SST*^+^ feedback *K* to zero.

Neurons receive external excitatory spike trains with Poisson statistics from two different sources. Both excitatory and *PV*^+^-interneurons, but not *SST*^+^-interneurons, receive feedforward input from the lateral geniculate nucleus in the thalamus (LGN) denoted by *J_s_* and where feedforward input to cortical *PV*^+^-interneurons is considerably stronger than the input to excitatory neurons by the factor *g_fw_*; this is consistent with experimentally measured inputs from the LGN to layer 4 (Ji et al., 2016), as well as from layer 4 to layer 2/3 (Pfeffer et al., 2013). Excitatory neurons and *SST*^+^-interneurons, but not *PV*^+^-interneurons, receive a second source of input denoted by *J_b_*, which we identified as background inputs from the surrounding cortical tissue or higher-order cortical centers (BKG) (Adesnik et al., 2012; Zhang et al., 2014). To minimize the number of parameters, we made these two inputs the same (Table 1). We can write the feedforward population connectivity vector as:

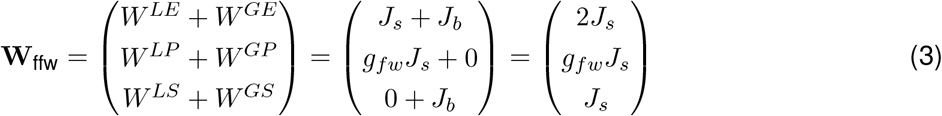

where *L* denotes LGN and *G* denotes background.

The total input current to neuron *i* from population *A* = {*E, P, S*}, *I^A^*(*t*), is the sum of synaptic input current from neurons in all populations *B* coupled to *A*, 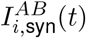, and the external input current, *I_i;ext_*(*t*):

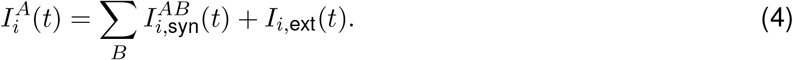

Denoting the spike train of neuron *j* in population *B* by 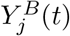 and the feedforward input spike train to neuron *i* by *X_i_*(*t*), we can further write:

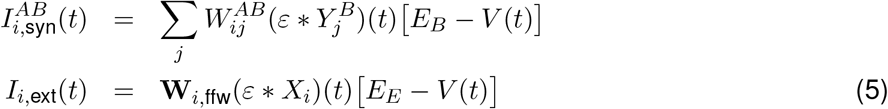

where *ε*(*t*) is an exponentially decaying synaptic kernel with a synaptic time constant *τ_syn_*, and * denotes a convolution. Note that here *E_E_* is the excitatory reversal potential, while EB could be either the excitatory reversal potential, *E_E_*, or the inhibitory reversal potential, *E_P_,E_S_*. To extract firing rates, we simulated the network for 30 seconds and calculated the time-averaged population rate. All network simulations were performed using NEST (Linssen et al., 2018).

### Implementation of deprivation-induced circuit changes

We introduced the parameters *δ_E_* < 1 and *δ_P_* < 1 to model the depression of feedforward synapses, and *ζ_EP_* > 1 and *ζ_PE_* > 1 to model the potentiation of recurrent synapses (Table 1). Synaptic changes induced by MD were applied multiplicatively to existing synaptic connections, with a new population connectivity matrix (Table 1):

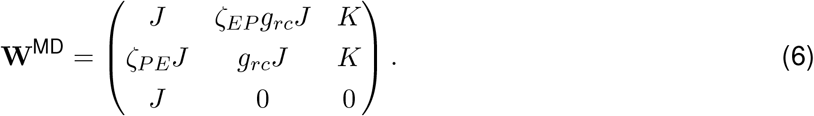

This is consistent with experimental measurements where the synaptic strength from excitatory to excitatory neurons and from fast-spiking to fast-spiking interneurons is unaffected, but other synapses potentiate (Maffei et al., 2006; Miska et al., 2018). There is no experimental evidence of plasticity in synapses of molecularly identified *SST*^+^-interneurons during MD. The new feedforward population connectivity vector after MD is:

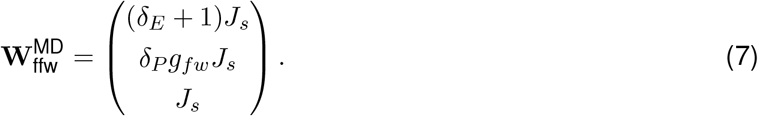

Note that we did not change the rate of the feedforward input, consistent with experimental findings (Linden et al., 2009).

To map firing rate changes in response to brief MD, we separately simulated networks with feedforward and recurrent MD-induced plasticity. For each type of synaptic change, the networks were simulated with parameter combinations on a 21 × 21 grid in the respective two-dimensional parameter planes (Figs. 1B, C and 4A, B). To study the interaction of feedforward and recurrent plasticity, we combined *δ_E_* and *δ_P_* into the E/I ratio of feedforward synaptic changes *ρ_EI_* = *δ_E_/δ_P_*. To simulate networks with varying E/I ratio, we fixed *δ_E_* = 1 and varied *δ_P_* (Figs. 1D and 4C) (see also (Miska et al., 2018) where *δ_E_* was also varied).

Feedforward plasticity in S1 was driven by a decrease of intrinsic excitability of *PV*^+^-interneurons through an increase of their firing threshold (Gainey et al., 2018). In the simulations, we thus changed the firing threshold of *PV*^+^-interneurons by adding *ξ_θ_*, without changing any feedforward inputs. For plasticity of the recurrent synaptic connections, we modeled the potentiation from *PV*^+^ to excitatory neurons (*ζ_EP_*) (Gainey et al., 2018). We simulated the S1 network with combined feedforward and recurrent plasticity following the procedure described above to extract the plane of firing rate changes (Fig. 6B, C).

### Quantification of the activity changes induced by deprivation

For each pair (*x,y*) of synaptic changes in the two-dimensional parameter space, (*x,y*) = (*δ_E_, δ_P_*) for feedforward plasticity, (*x,y*) = (*ζ_EP_,ζ_PE_*) for recurrent plasticity and (*x,y*) = (*ζ_PE_,ρ_EI_*) for combined feedforward and recurrent plasticity, we obtained a matrix of rate fold-changes of population *A* = {*E,P, S*} at MD, 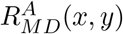, relative to baseline, 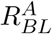:

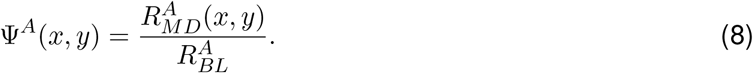

To quantify the responses, we first studied the fractional area of facilitation. Elements of Ψ^*A*^(*x,y*) were thresholded at 1, which denotes the border between suppression (Ψ^*A*^(*x,y*) < 1) and facilitation (Ψ^*A*^(*x,y*) > 1). All matrix elements corresponding to facilitation were then set to 1, while those for suppression to −1. Summing up all the positive entries in this thresholded matrix and dividing by the total number of matrix elements produced the fractional area of facilitation (*F_E_, F_P_* in Figs. 2 and 4). We calculated the overlap of the response areas in excitatory neurons and *PV*^+^-interneurons by multiplying the thresholded matrices of excitatory and *PV*^+^ responses element-wise. After this multiplication, matrix elements corresponding to pairs of synaptic changes that resulted in the simultaneous suppression or facilitation of excitatory and *PV*^+^ responses had values of +1, while elements corresponding to opposite responses of excitatory and *PV*^+^ had values of −1. We summed all positive values and divided this sum by the total number of matrix elements to calculate the fractional area of the overlap for facilitation and suppression in the two neural populations (*O_EP_* in Figs. 2 and 4).

### Capturing the sequence of firing rate suppression and recovery

We extracted the experimental data for firing rate fold changes of excitatory and fast-spiking interneurons during the first two days of MD from Figure 3D in (Hengen et al., 2013) using open source software (Rohatgi, 2011). We hypothesized that MD should affect feedforward connections immediately. Therefore, to capture the firing rates of excitatory and *PV*^+^ neurons on the first day of MD, we assumed that only feedforward synapses change (Fig. 7, left). We simulated networks in the ISN regime with strong *SST*^+^-feedback for 70 randomly drawn (*δ_E_, δ_P_*)-pairs chosen from a range in which excitatory neurons have normalized firing rates close to baseline (1.05 ± 0.15 in Fig. 4A). This set of parameters produces suppressed *PV*^+^ firing rates.

To capture the firing rates of excitatory and *PV*^+^ neurons on the second day of MD, we assumed that in addition to changes in the feedforward connections, recurrent connections also change (Fig. 7, right). Therefore, for each (*δ_E_,δ_P_*)-pair, we found a (*ζ_EP_,ζ_PE_*)-pair that can recover the firing rates of *PV*^+^-interneurons close to baseline (0.98 ± 0.15 in Fig. 4B).

We also simulated the ISN without *SST*^+^-feedback with the same changes in feedforward and recurrent connections to investigate how the firing rate changes in this network compare with those measured experimentally.

### Linear population-rate model

To gain a deeper understanding of the factors underlying firing rate changes in response to MD-induced plasticity, we used a Wilson-Cowan linear population model where we studied the rates of each population (Wilson and Cowan, 1972). The dynamics of the firing rate in this model follow:

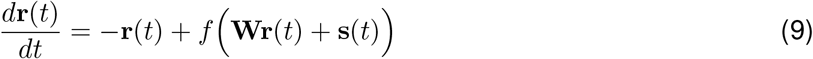

where **r** is the vector of rates, s the vector of external inputs and **W** the population connectivity matrix of the rate-based model:

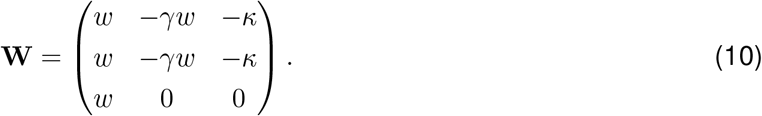

As in the simulated network, input to the network comes from two sources, LGN and BKG. As before, we set the input rates from both sources to be the same, *r_x_*, in which we also absorb the input weights:

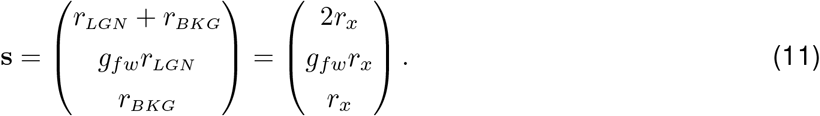

We used a rectified-linear population input-output function:

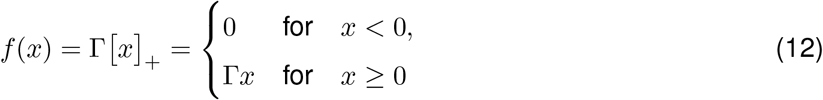

where Γ is the input-output gain, chosen to be the same for all populations. With steady inputs and assuming that the network operates in the linear regime away from rectification, the steady-state of the rates in this model is:

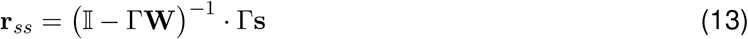

From this steady-state, we can calculate the fold-change of the network rate after MD relative to baseline for each combination of synaptic changes. Synaptic changes induced by MD were applied multiplicatively as before, with (*δ_E_,δ_P_*) and (*ζ_EP_,ζ_PE_*) affecting **W** and s, similarly to Eqs. 6 and 7, to yield:

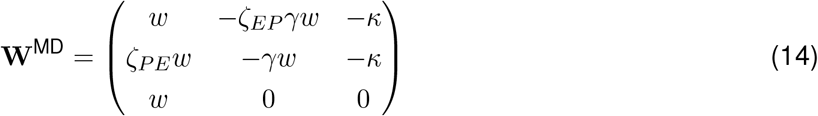

and

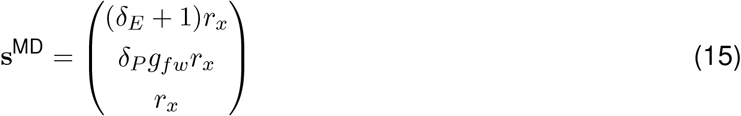

We solve the steady state in Eq. 13 for MD by inverting 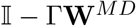. Without loss of generality, we absorb the response gain factor Γ into the interaction parameters (*w* and *κ* respectively) as well as into the feedforward input (*r_x_*) by rescaling them with a common factor. The steady-state activity after MD becomes:

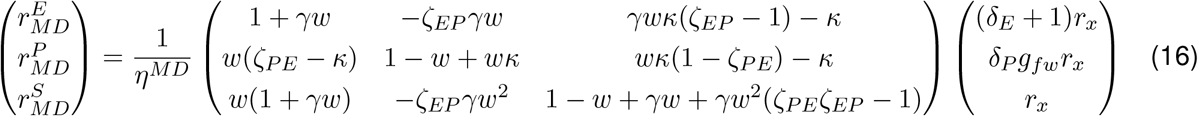

where

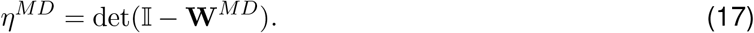

To generate the planes of firing rate change in response to MD-induced plasticity, we solved the rate model in Eq. 9 numerically, while also incorporating firing rate rectification when rates became negative. We found that the rate model could capture firing rates changes in the spiking networks both for the network with a single subtype of interneuron (compare Supplementary Fig. S2 A-C vs. Fig. 1 B-D and Supplementary Fig. S2 D-F vs. Fig. 2 B-D) and for the network with two subtypes of interneurons (compare Supplementary Fig. S3 vs. Fig. 4).

### Analysis of the paradoxical effect and its reversal

The linear model enables us to mathematically relate the firing rate changes to MD-induced plasticity to specific response properties of the network depending on its operating regime. In the linear model, setting the coupling scale *w* = 1 marks the transition between the non-ISN (*w* < 1) and the ISN (*w* > 1) regimes (Tsodyks et al., 1997). A hallmark of the ISN regime is the “paradoxical effect” whereby an excitatory input to the inhibitory population decreases its firing rate after a transient increase.

We first considered the network with a single subtype of interneuron, by setting *κ* = 0. In this case the firing rate change in response to additional drive > 0 to the *PV*^+^ population becomes:

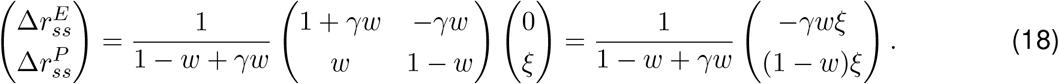

The scalar pre-term 1 – *w* + *γw* is always positive when the network is stable. The responses of each population to this additional drive to the *PV*^+^ population are:

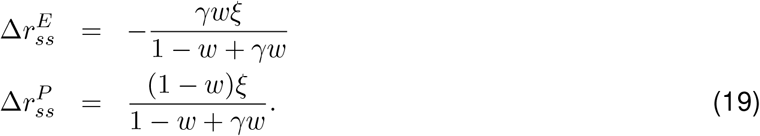

Hence, the response of the excitatory neurons is always in the opposite direction to for any network parameters (*γ* > 0 and *w* > 0). However, the response of the *PV*^+^ population depends on the value of *w*. When *w* < 1 and the network is in the non-ISN regime, the *PV*^+^ population changes its activity in the same direction as ξ. When *w* > 1 and the network is in the ISN regime, the *PV*^+^ population changes its activity in the opposite direction to ξ, as the excitatory population. Since the depression of synaptic connections to *PV*^+^-interneurons observed in early MD is equivalent to providing an inhibitory input to the *PV*^+^ population in the linear model, i.e. *ξ* < 0, the operating regime of the network and the presence of the paradoxical effect in the ISN regime can be related to the observed firing rate changes in response to MD. Therefore, in the non-ISN, the excitatory and *PV*^+^ populations respond to MD by changing their firing rates in the opposite direction, while in the ISN regime, the excitatory and *PV*^+^ firing rates changes closely follow each other (Supplementary Fig. S2). We confirmed through simulation that the value of *J* where the same change in *PV*^+^ responses to MD-induced plasticity occurs in the spiking network is indeed the point where the spiking network goes from non-ISN to ISN and is related to the paradoxical effect (Fig. 2).

We extended this result in the network with two types of interneurons. Now the firing rate change in response to additional drive > 0 to the *PV*^+^ population becomes:

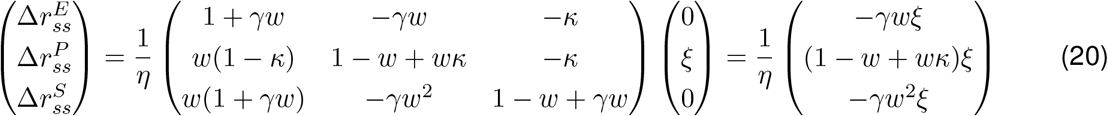

where 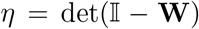, which is always positive when the network is stable. The responses of the excitatory and the *PV*^+^ population to this additional drive to the *PV*^+^ population are:

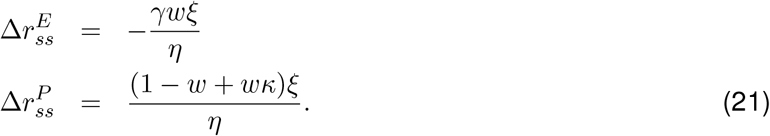

As for the network with a single type of interneurons, the response of the excitatory neurons is always in the opposite direction to *ξ* for any network parameters, independent of the strength of *SST*^+^ feedback *κ*. The response of the *PV*^+^ population is proportional to (1 – *w* + *wκ*)*ξ*. Thus, the direction of rate changes in the *PV*^+^ population no longer depends only on *w* but also on the strength of *SST*^+^ feedback *κ*. In the non-ISN, the *PV*^+^ population responds with a change of activity in the same direction as *ξ* since *w* < 1 and *wκ* > 0, independent of *κ*. In the ISN, where *w* > 1, the *PV*^+^ population responds with a change of activity in the opposite direction to when the following condition is satisfied:

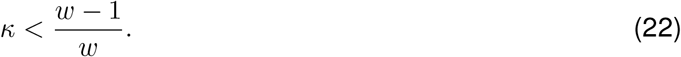

This implies that for any *w* > 1, there exists a sufficiently large *κ* that leads to a reversal of the paradoxical effect and thus a non-paradoxical response of the *PV*^+^ population even in a network operating as ISN (Fig. 5).

### Relating facilitation of *PV*^+^ firing rate to the paradoxical effect

Next, we related the above conditions (Eq. 22 and *w* > 1) for the presence of the paradoxical effect in networks with one or two types of interneurons to the presence of facilitatory responses for *PV*^+^-interneurons in the parameter spaces of MD-induced plasticity. In particular, in the network with a single type of interneuron, *PV*^+^-interneurons start showing a facilitatory response in response to feedforward plasticity at a specific value of coupling (Supplementary Fig. S2D). In the network with two subtypes of interneurons operating in the ISN regime, the facilitatory response of *PV*^+^-interneurons vanishes for a specific strength of *SST*^+^ feedback *κ* and reemerges at a higher value of *κ* (Fig. S3D).

From Eq. 16, the rate for the *PV*^+^ population after induction of MD-plasticity is:

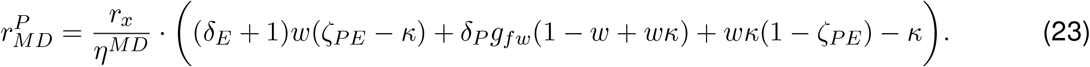

We focus on feedforward plasticity only, since recurrent plasticity affects *η^MD^* (Eq. 17). In the case of feedforward plasticity, the rate for the *PV*^+^ population after induction of MD-plasticity reduces to:

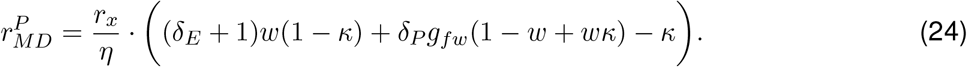

Setting all MD-related parameters to one, the baseline rate is:

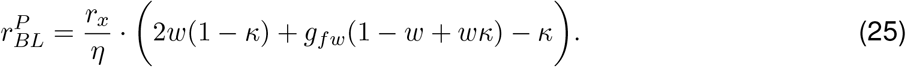

Therefore, the condition for the emergence a facilitatory area of *PV*^+^ responses after MD is:

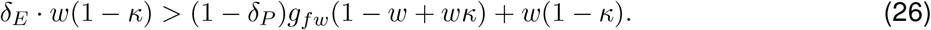

We first considered the network with a single type of interneuron by setting *κ* = 0. Then the condition for the emergence a facilitatory area of *PV*^+^ responses after MD becomes:

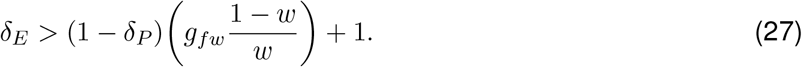

Since feedforward plasticity in response to MD depresses synaptic inputs to excitatory neurons and *PV*^+^-interneurons, *δ_E_, δ_P_* < 1. Therefore, this condition can never be satisfied in the non-ISN regime where *w* < 1. Thus, *PV*^+^ responses to MD-induced feedforward depression in the non-ISN never increase above baseline (Supplementary Fig. S2D).

In the ISN regime, *w* > 1, the condition becomes:

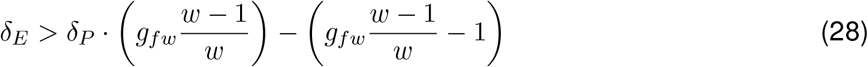

giving a specific linear relationship between *δ_E_* and *δ_P_* that is the boundary between suppression and facilitation (Supplementary Fig. S2A and 1B). Thus, the position of the boundary between facilitation and suppression depends on *w* and *g_fw_*. However, for *w* » 1, which is well in the ISN regime, (*w* – 1)/*w* ≈ 1 and thus the slope and offset of this linear relationship depend primarily on *g_fw_*.

In the ISN network with *SST*^+^-interneurons where *w* > 1, we studied the condition for the emergence a facilitatory area of *PV*^+^ responses for increasing *κ*. Assuming that 0 < *κ* < 1, the condition yields again a linear relation between *δ_E_* and *δ_P_* that separates facilitatory and suppressive areas of the parameter space:

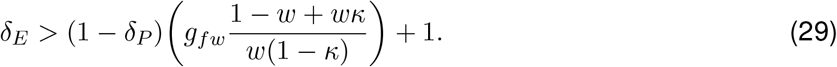

As long as

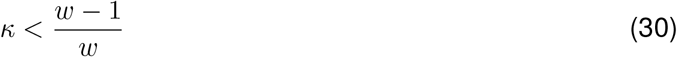

the condition Eq. 29 can be satisfied with depression of feedforward synapses. Note that this is exactly the condition Eq. 22 for which the *PV*^+^ population switches to non-paradoxical behavior due to *SST*^+^ feedback. As *κ* increases, the condition Eq. 29 becomes harder to satisfy because the term 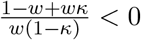 grows monotonically. Thus, the facilitation area of the *PV*^+^ population in the parameter space of feedforward plasticity decreases (Fig. S3D). As *κ* increases, the condition Eq. 29 can no longer be satisfied, and *PV*^+^-responses no longer show facilitation (Fig. S3D; at *κ* = 0.8 for the chosen parameters, see also switch of paradoxical response in Fig. 5).

As *κ* approaches one, the condition in Eq. 26 goes through a singularity. Therefore, the condition for the emergence of facilitation of *PV*^+^ responses after MD in Eq. 29 switches sign:

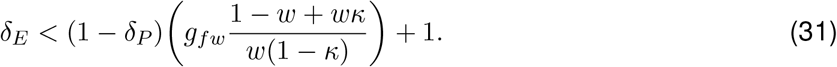

*PV*^+^-interneurons can again show a facilitatory response during MD-induced plasticity for *κ* > 1, but on the opposite side of the linear boundary (Fig. S3A).

For networks well in the ISN regime, where *w* > 1, the conditions for the disappearance of the facilitatory area and the reemergence on the opposite side of the linear boundary become the same because (*w* – 1)/*w* → 1. Here, the condition for reversal of the paradoxical effect in *PV*^+^-interneurons through *SST*^+^ feedback and for the opposite firing rate changes of excitatory neurons and *PV*^+^-interneurons through *SST*^+^ feedback become the same.

## Supplementary figures

**Supplementary Figure S1:**
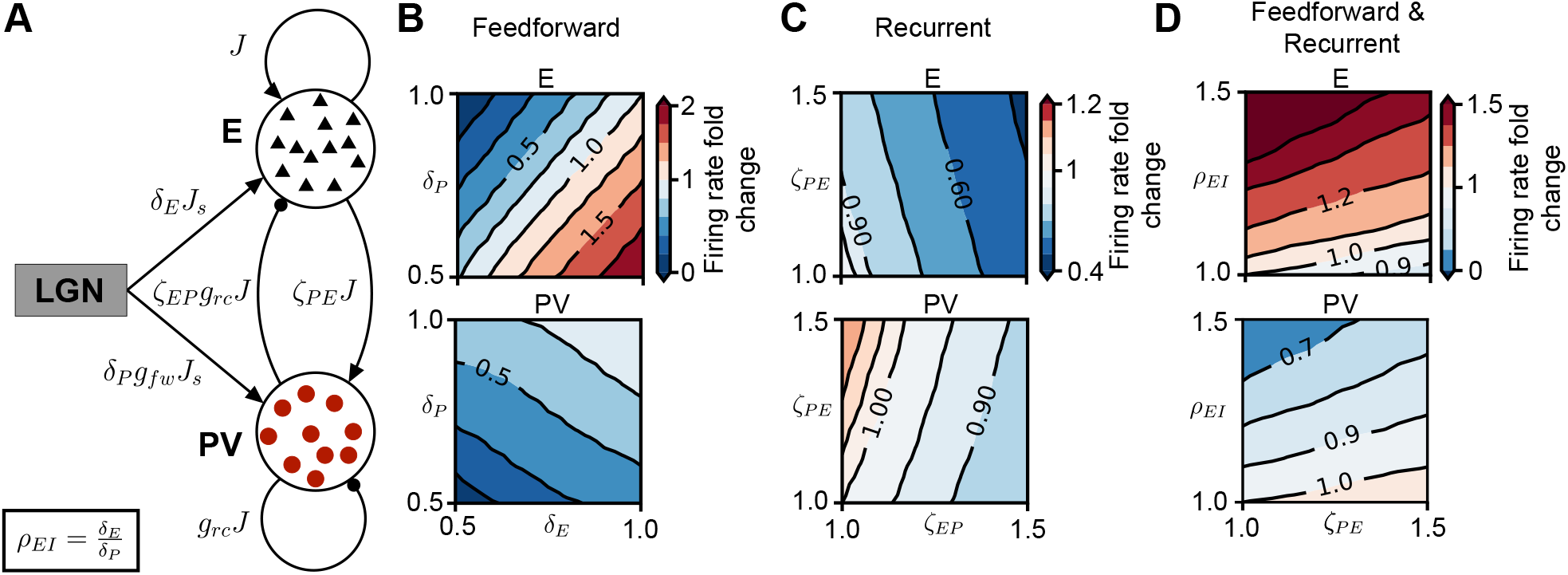
Response to synaptic changes induced by brief MD in a spiking network in non-ISN regime. **(A)** Network schematic showing couplings among cells and parameters to model MD-induced synaptic changes (see Fig. 1). In the non-ISN, *J* has a smaller value than in the ISN (shown responses for *J* = 0.01 nS) **(B)** Network firing rate in the (*δ_E_, δ_P_*) plane as fold-change of baseline firing rate for excitatory neurons (top) and *PV*^+^-interneurons (bottom). In the non-ISN there is no facilitation of activity of *PV*^+^. **(C)** Network firing rate in the (*ζ_EP_, ζ_PE_*) plane as fold-change of baseline firing rate. Facilitation of activity of *PV*^+^-interneurons is present in the non-ISN. **(D)** Combined feedforward (through the E/I ratio of feedforward synaptic changes, *ρ_EI_* = *δ_E_/δ_P_*) and recurrent plasticity (through the potentiation of recurrent excitation to *PV*^+^ -interneurons, *ζ_PE_*). Network firing rate as fold-change of baseline firing rate in (*ζ_PE_*, *ρ_EI_*) plane. In contrast to the ISN (see Fig. 1), here in the non-ISN, the rate changes are opposite over large fractions of the plane.

**Supplementary Figure S2:**
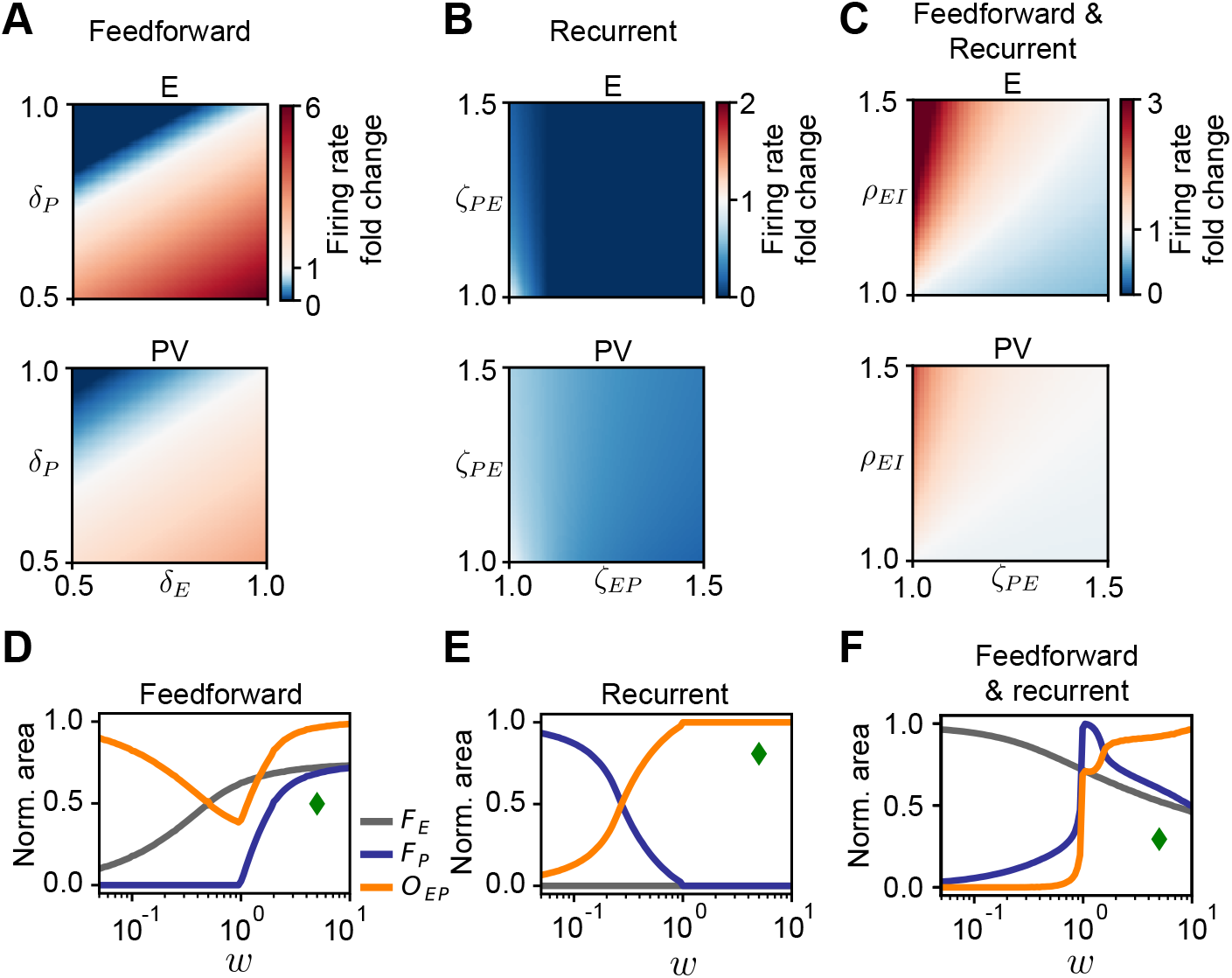
Linear model of the network with a single subtype of interneurons (*PV*^+^). **(A)** Network firing rate in the (*δ_E_, δ_P_*) plane as fold-change of baseline firing rate for excitatory (top) and *PV*^+^ populations (bottom). **(B)** Network firing rate in the *(ζ_EP_,ζ_PE_*) plane as fold-change of baseline firing rate. **(C)** Combined feedforward (through the E/I ratio of feedforward synaptic changes, *ρ_EI_* = *δ_E_/δ_P_*) and recurrent plasticity (through the potentiation of recurrent excitation to *PV*^+^ -interneurons, *ζ_PE_*). Network firing rate as fold-change of baseline firing rate in *(ζ_PE_, ρ_EI_*) plane. **(D)** Network with feedforward depression only. Fractional area of facilitation for excitatory neurons (*F_E_*, gray), *PV*^+^-interneurons (*F_P_*, blue) and the overlap between excitatory and *PV*^+^ response areas (*O_EP_*, orange) as a function of overall coupling scale *w*. **(E)** Same as (D) for recurrent potentiation only. **(F)** Same as (D) for combined feedforward and recurrent plasticity. For all plasticity types studied, the linear model qualitatively reproduces the structure of firing-rate fold changes in the spiking network (compare to Fig. 1 and Fig. 2). Note that w in the linear population model is the equivalent to the coupling scale *J* in the spiking network. Green diamond shows the coupling scale value (*w* = 5) used in (A-C).

**Supplementary Figure S3:**
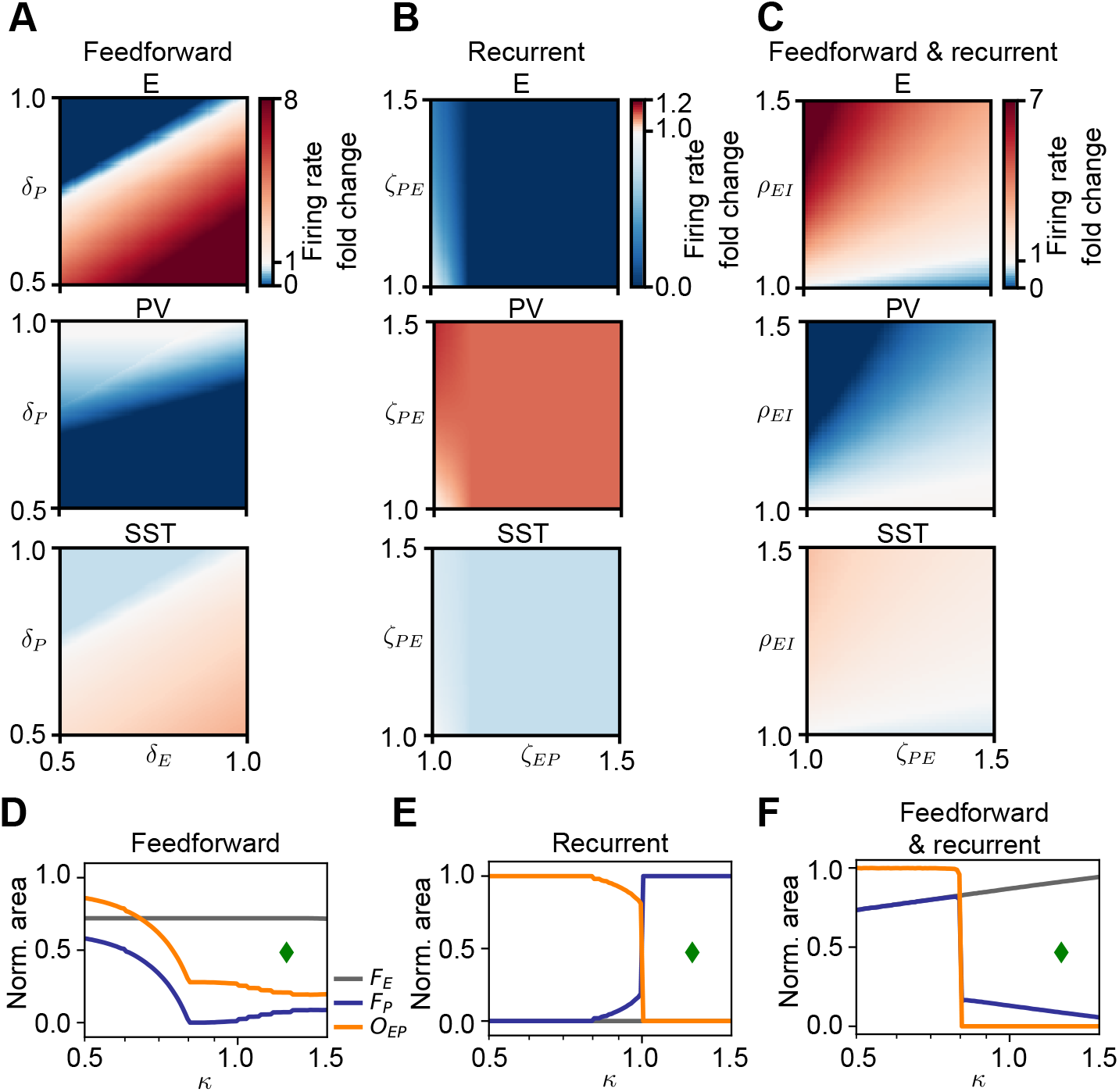
Linear model of the network with a two subtypes of interneurons (*PV*^+^ and *SST*^+^). **(A)** Population rate in the (*δ_E_,δ_P_*) plane as fold-change of baseline in the linear model containing *SST*^+^-interneurons. **(B)** Population rate in the (*ζ_EP_,ζ_PE_*) plane as fold-change of baseline in the linear model containing *SST*^+^-interneurons. **(C)** Population rate in the (*ζ_EP_,ρ_EI_*) plane as fold-change of baseline in the linear model containing *SST*^+^-interneurons. **(D-F)** Fractional area of facilitation for excitatory neurons (*F_E_*, gray), *PV*^+^-interneurons (*F_P_*, blue) and the overlap between excitatory and *PV*^+^ response areas (*O_EP_*, orange) as a function of *SST*^+^ feedback κ, for the three types of plasticity considered: feedforward only (D), recurrent only (E), and combined feedforward and recurrent (F). Green diamond shows the value of *κ* used in (A-C) for the response planes showing reversed responses in *PV*^+^-interneurons.

**Supplementary Figure S4:**
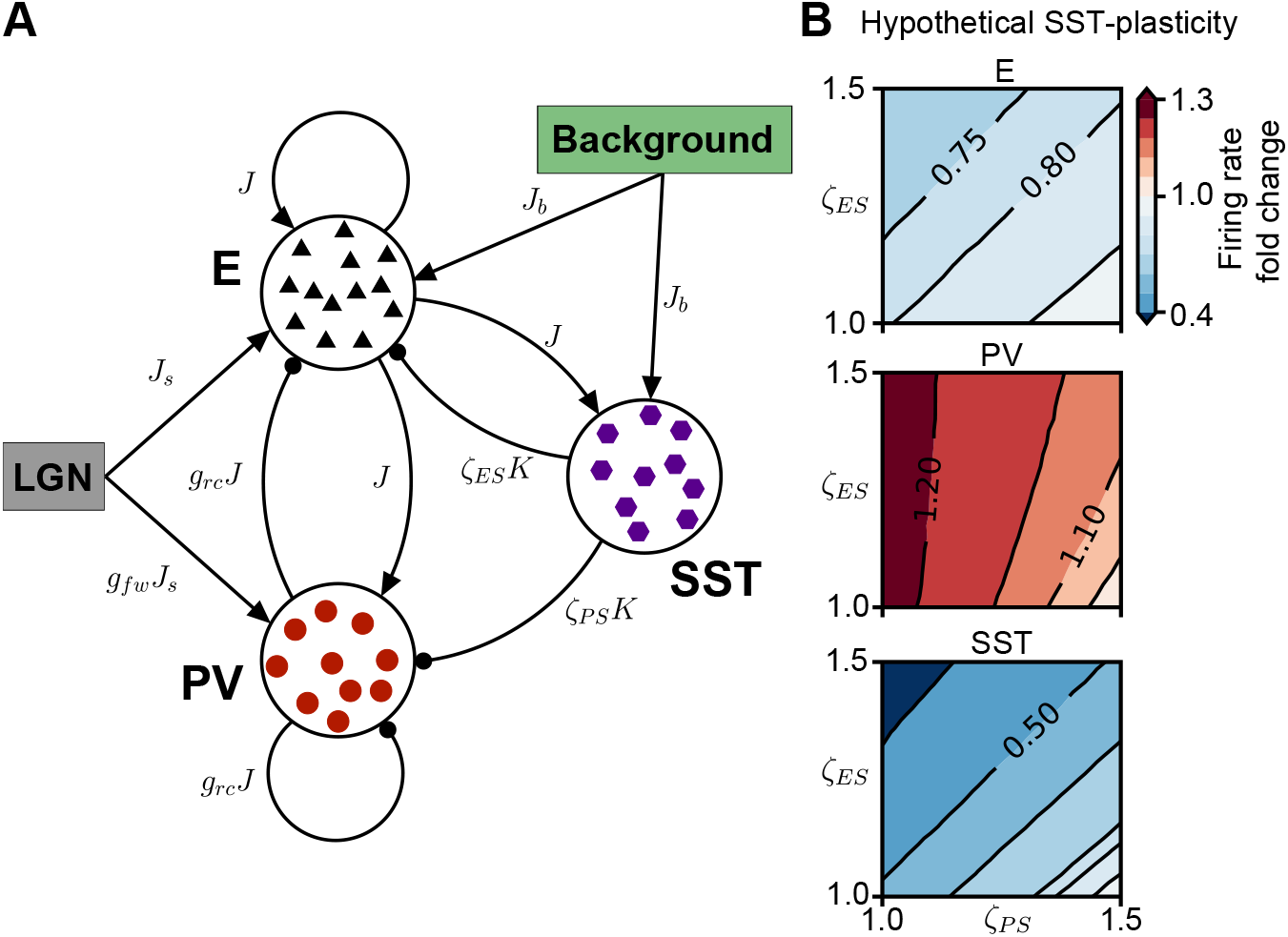
Plasticity of synapses from *SST*^+^-interneurons affects circuit dynamics similarly to potentiation of recurrent synapses between excitatory and *PV*^+^-interneurons. **(A)** Network schematic showing couplings among cells and parameters to model hypothetical MD-induced synaptic changes in synapses from *SST*^+^-interneurons: depression of synapses from *SST*^+^-onto *PV*^+^-interneurons *(ζ_PS_* < 1) and potentiation of synapses from *SST*^+^-interneurons onto excitatory neurons *(ζ_ES_* > 1). **(B)** Network firing rate in the *(ζ_PS_,ζ_ES_*) plane as fold-change of baseline firing rate (bottom right corner where *ζ_PS_* = *ζ_ES_* = 1). The qualitative effect of this hypothetical plasticity closely resembles that of recurrent plasticity (Fig. 4B).

## Notes

### Competing Interest Statement

The authors have declared no competing interest.

